# Morphometry-based detection of deep learning faults in glomerular segmentation

**DOI:** 10.64898/2025.12.23.696291

**Authors:** Hrafn Weishaupt, Justinas Besusparis, Nazanin Mola, Ståle Sund, Sabine Leh

## Abstract

Deep learning-based segmentation has evolved to a powerful strategy for automatically annotating glomeruli in kidney biopsy images. However, since any artificial intelligence can make mistakes, strategies for identifying and correcting faulty annotations are often indispensable. Yet, how can such a validation be achieved without the laborious task of a pathologist manually checking every single image? To address this issue, the current project performed an extensive study on the use of shape analysis to automatically evaluate the glomerular annotations produced by deep-learning segmentation. Examining a large repertoire of shape descriptors on over 168000 glomerular predictions, the study found that morphometry could successfully highlight and distinguish between three different types of segmentation inconsistencies. In addition, using shape descriptors to rank segmentation annotations, it was possible to obtain a distinct enrichment of errors on the leading edge of the ranking, implying that pathologists would only have to inspect and correct the most suspicious fraction of all annotations. Ultimately, the study suggested a panel of three shape descriptors that enabled an efficient enrichment of all errors, respective or irrespective of error type. In summary, the work demonstrates the methodological aspects and benefits of shape analysis for evaluating glomerular segmentation results. We are convinced that, by applying such a strategy for detecting segmentation errors, it will be possible to approach a more time-efficient correction of deep learning-derived glomerular annotations.

## 1 Introduction

With the advent of digital pathology and the associated accessibility of digitized biopsy images, deep learning (DL) has emerged as a promising strategy for aiding pathologists in various diagnostic tasks [1]. For instance, in nephropathology, one of the most important tasks in evaluating a kidney biopsy is the characterization of morphological lesions in glomeruli [2], clusters of capillaries responsible for the filtration of blood and production of urine. In this context, deep learning has shown great progress in both automatically detecting/segmenting such structures in whole slide images (WSIs) [3–11], as well as classifying them into defined lesion categories [11–17].

While few if any of such DL models have yet entered clinical use in any validated diagnostic workflow, they still present a highly valuable resource for enhancing ongoing research. For instance, the automatic detection/segmentation of glomeruli can aid tremendously in the generation of large-scale collections of glomerular image patches, in turn easing the development and training of downstream models for e.g. classification [13, 16, 17]. However, when using DL for the gathering of training data, segmentation mistakes [3, 8, 9] could have adverse consequences and would often need to be detected and corrected before using the data for downstream purposes. Nevertheless, manually screening for such errors then represents again a time-consuming obstacle towards the automatic creation of glomerular image collections (Fig. 1A-C).

**Fig. 1.**
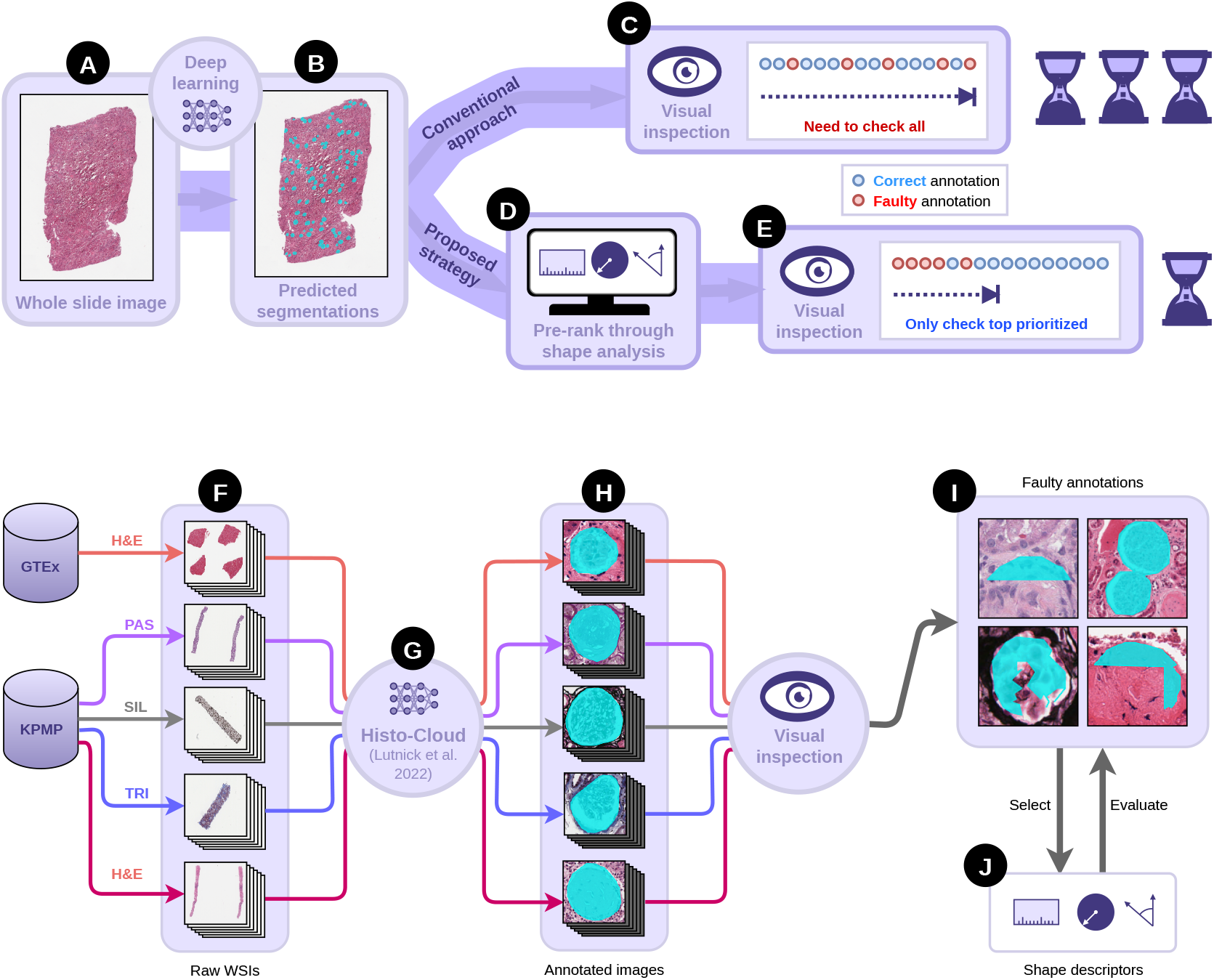
Illustration of goal (upper panel, A-E) and overall workflow (lower panel, F-J). Upper panel: To mitigate the time-consuming task of manually checking all annotations after an automatic glomerular segmentation (A-C), the project aims to identify a set of metrics (D) via which annotations with putative faults can be prioritized during inspection (E). Lower panel: Subjecting whole slide images (WSIs, F) from the Genotype-Tissue Expression (GTEx) and the Kidney Precision Medicine Project (KPMP) to the Histo-Cloud tool [4] for automatic segmentation (G), a large repository of candidate glomerular patches is extracted (H). Annotations are visually inspected to identify categories of segmentation faults (I), and the faults are used to guide the selection and evaluation of metrics for automatically prioritizing such errors (J).

Thus, in order to make full use of available DL models for generating research data sets, open questions that need to be addressed are: “How can we estimate the number of mistakes in the absence of a ground truth?”, and “how can we efficiently detect and correct mistakes, without requiring a pathologist to manually revisit each and every image?” For instance, would it be possible to devise a set of automatic analysis metrics that, when applied to a collection of segmentation annotations, would strongly enrich for errors, thus allowing the pathologist to focus corrections to the most suspicious cases (Fig. 1D-E)?

Considering that the instance segmentation of glomeruli from kidney biopsy images produces annotations that can be interpreted as two-dimensional shapes, i.e. polygons, morphometry/shape analysis lends itself as a promising strategy to identify inconsistent segmentations. Shape analysis denotes the quantitative and objective investigation of shape features and has broad applications in the study of biomedical image objects, including e.g. biological cells [18, 19], tumors [20, 21], calcifications in mammograms [22], and even glomeruli [23, 24]. More importantly, shape analysis can be employed to evaluate the results of image segmentations [8, 25], or as a means to help with the refinement/correction of initial segmentation results [8, 9]. For instance, with respect to glomerular segmentation, Altini et al. [8] used the difference between an annotation’s area and the area of its convex hull as a measure to identify annotations that might harbor multiple glomeruli. Trinh et al. [9] suggested a post-processing step converting each annotation to its convex hull to remove holes and irregularities. In line with these studies, we recently also reported the use of circularity and convexity to filter glomerular segmentation predictions [26]. However, very little work has yet focused on a more extensive investigation into which types of annotation errors can be expected from deep learning-based glomerular segmentation or how to best describe and detect such errors using shape analysis.

To the best of our knowledge, the current work describes the first more systematic study on shape analysis for detecting annotation errors in glomerular segmentation. Specifically, the project seeks to (i) reveal the common types of segmentation mistakes that can appear in large scale automatically segmented glomerular collections and to (ii) evaluate a broad panel of shape descriptors in their ability to automatically highlight such errors.

## 2 Approach

To evaluate the use of shape analysis as a tool to prioritize glomerular segmentations in need of correction, the study pursued a strategy as outlined in figure 1F-J. Specifically, utilizing a diverse collection of WSIs and applying an existing DL tool for glomerular segmentation, the study obtained a large number of predicted glomerular annotations. Subjecting these annotations to visual inspection, the various types of segmentation errors were identified. Subsequently, various shape descriptors were selected or designed to detect such errors, and their usability was evaluated through qualitative and quantitative analyses. The individual steps in this procedure are briefly described below. For a more in-depth description of the individual data sets and methods, the reader is referred to the Supplementary Material.

## 3 Material and methods

### 3.1 Collection of raw WSIs

To obtain a sufficiently large and diverse collection of WSIs across different tissue contents and histological stains, the project acquired raw WSIs from two public resources (Fig. 1F): the Kidney Precision Medicine Project (KPMP)[27], including biopsies stained with hematoxylin & eosin (H&E, n=500), periodic acid-Schiff (PAS, n=500), Jones’ methenamine silver (SIL, n=500), and trichrome (TRI, n=500), and the Genotype-Tissue Expression (GTEx, n=500) project [28], which contains autopsy tissues stained with H&E.

### 3.2 Automatic segmentation of glomeruli

The collected WSIs were subjected to an automatic DL-based instance segmentation via the Histo-Cloud tool [4](Fig. 1G), as also described in our previous studies [26, 29–31], producing a JSON file with a polygon-shaped annotation for each detected object (Fig. 1H) in an image. The final collection comprised 171991 annotations, spanning 105646 objects extracted from GTEx WSIs, and 66345 objects (H&E: 12407; PAS: 18102; SIL: 18311; TRI: 17525) extracted from KPMP WSIs.

### 3.3 Detection of invalid annotations

In some cases, an erroneous segmentation could be directly identified by determining whether the corresponding annotation represented an invalid polygon, i.e. a polygon with intersecting boundary lines or intersections at boundary points (Supp. Fig. 1). As such mistakes could be detected without requiring any shape analysis, they were filtered out prior to any subsequent analysis (*n* = 3252). In addition, some annotations (*n* = 8) were clear outliers just by considering their area (> 200000 µm^2^), and were also removed prior to downstream analyses. After filtering, 168731 annotations were left for downstream analyses.

### 3.4 Characterization of segmentation errors

Following the segmentation and removal of invalid polygons, a markup image patch was extracted for each annotation (Fig. 1H). Subsequently, the patches were visually screened to discover predominant types of segmentation faults (Fig. 1I), revealing five broad categories of errors, i.e. incomplete annotations with (i) a vertical or horizontal cut (Fig. 2A-B, Supp. fig. 2A-B), (ii) a diagonal cut (Fig. 2C-D, Supp. fig. 2C), or (iii) an irregular/rough boundary (Fig. 2E-F, Supp. fig. 2D), (iv) single annotations including multiple glomeruli (Fig. 2G-H, Supp. fig. 2E), and (v) false-positive detections (Fig. 2I-J). In addition, some glomeruli were missed during segmentation, accounting for false negatives (Supp. fig. 3C-D). From hereon, categories i-ii are referred to as *cut*, category iii is referred to as *irregular*, category iv is referred to as *multi-object*, and false positives and false negatives are referred to as *FP* and *FN*, respectively.

**Fig. 2.**
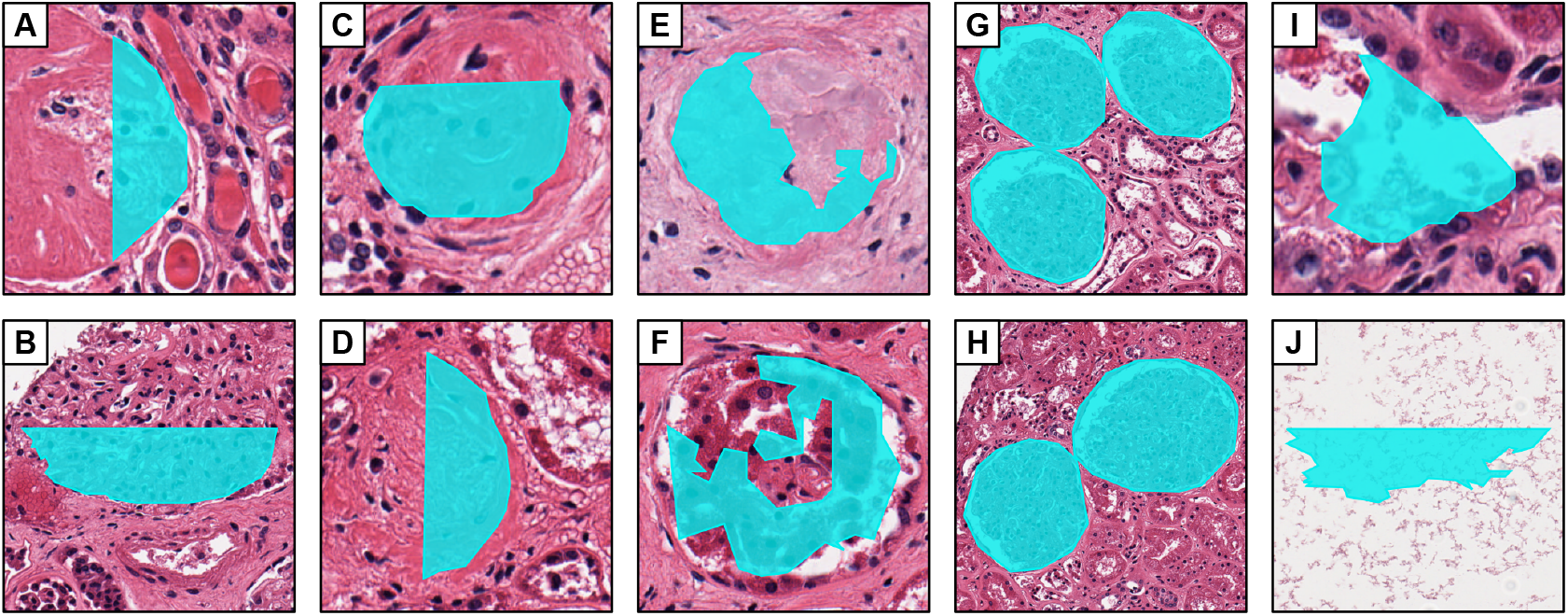
Examples of segmentation errors observed in the GTEx WSIs. **A-F)** Incomplete segmentation of glomeruli resulting either in regular shaped annotations with a clear cut (A-D) or irregular shaped annotations (E-F). The cut in A and B is vertical or horizontal, respectively, suggesting inconsistencies between adjacent tiling windows, while the cuts in C and D are diagonal but nearly vertical or horizontal. **G-H)** Segmentation of multiple adjacent glomeruli as a single object. **I-J)** False-positive detections.

Subsequently, the frequency of each of these error types was estimated through manual human labeling on a random subset of segmentation objects (counting cut, irregular, multi-object, FP annotations) or WSIs (counting FNs), respectively.

### 3.5 Selection of shape descriptors

Having determined the possible types of faulty annotations, a set of metrics was selected and evaluated for automatically detecting specifically the cut, irregular, and multi-object annotations (Fig. 1J). Towards this end, the study explored a selection of geometric descriptors designed to capture aspects of the general shape and of boundary irregularity. Following DL segmentation, each predicted glomerulus (Fig. 3A) was represented by an annotation in form of a polygon (PG; Fig. 3B). Adding additional variations of the annotation (Fig. 3C) in terms of the convex hull (CH), minimum bounding box (BB), and minimum rotated bounding box (BBrot), various geometric quantities were computed (Fig. 3D, Table 1), and then utilized to calculate a set of shape descriptors (Table 2).

**Fig. 3.**
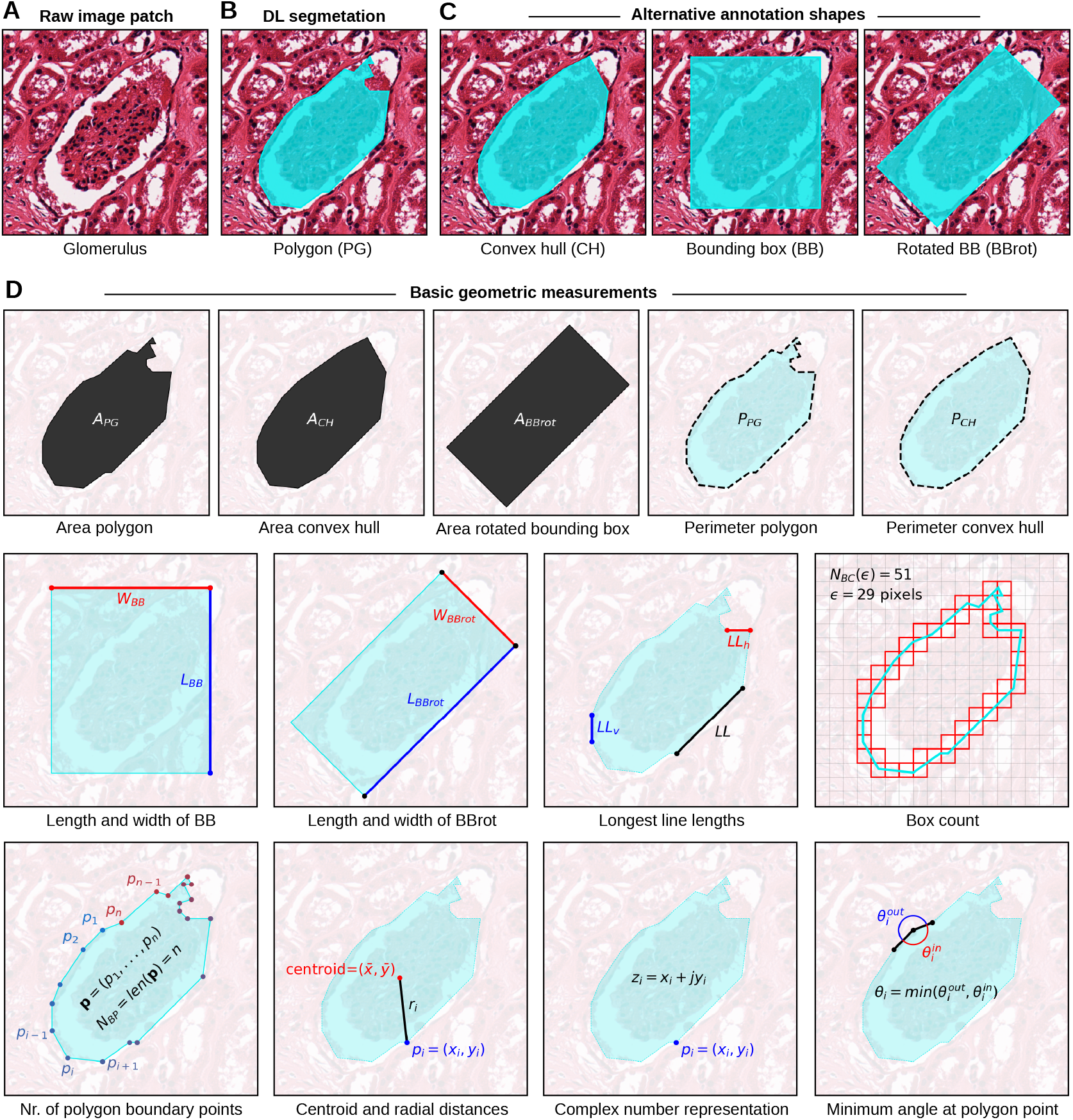
Schematic overview over the various basic geometric shapes/properties extracted for each annotation. **A)** Image patch with a full glomerulus in the center. **B)** Polygon (PG) annotation obtained from segmentation. **C)** Three alternative geometric shapes used to annotate the glomerulus: Convex Hull (CH; left), minimum Bounding Box (BB; center), and minimum rotated Bounding Box (BBrot; right). **D)**. Basic geometric properties, including areas and perimeters, bounding box dimensions, the longest boundary line segments (general, horizontal, or vertical), the number of grid boxes intersected by the boundary line, number of boundary points, the distances of points to the centroid of the polygon, the complex number representation of points, and the angle between two adjacent boundary line segments.

**Table 1.**
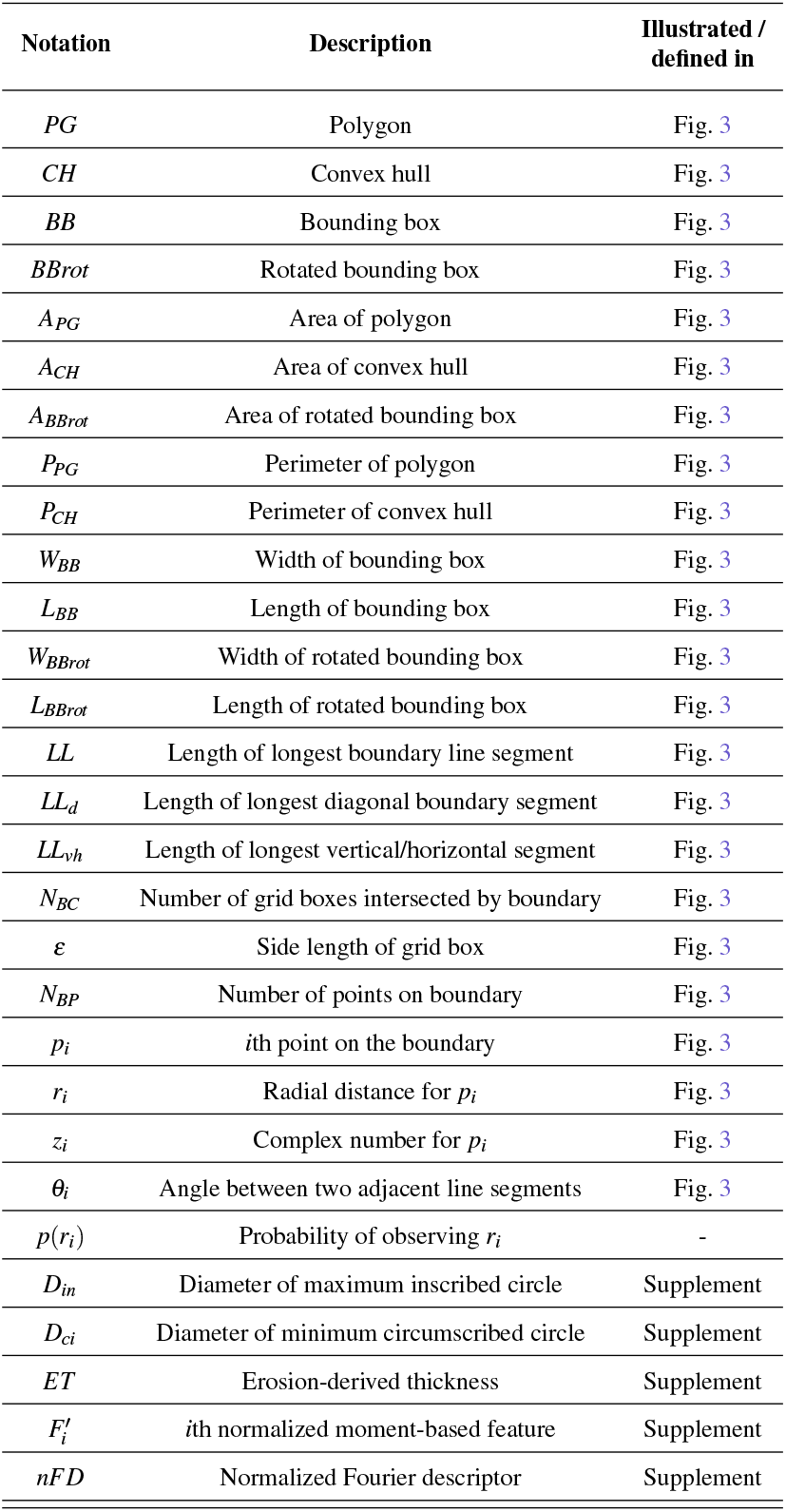
List of base geometric notations.

**Table 2.**
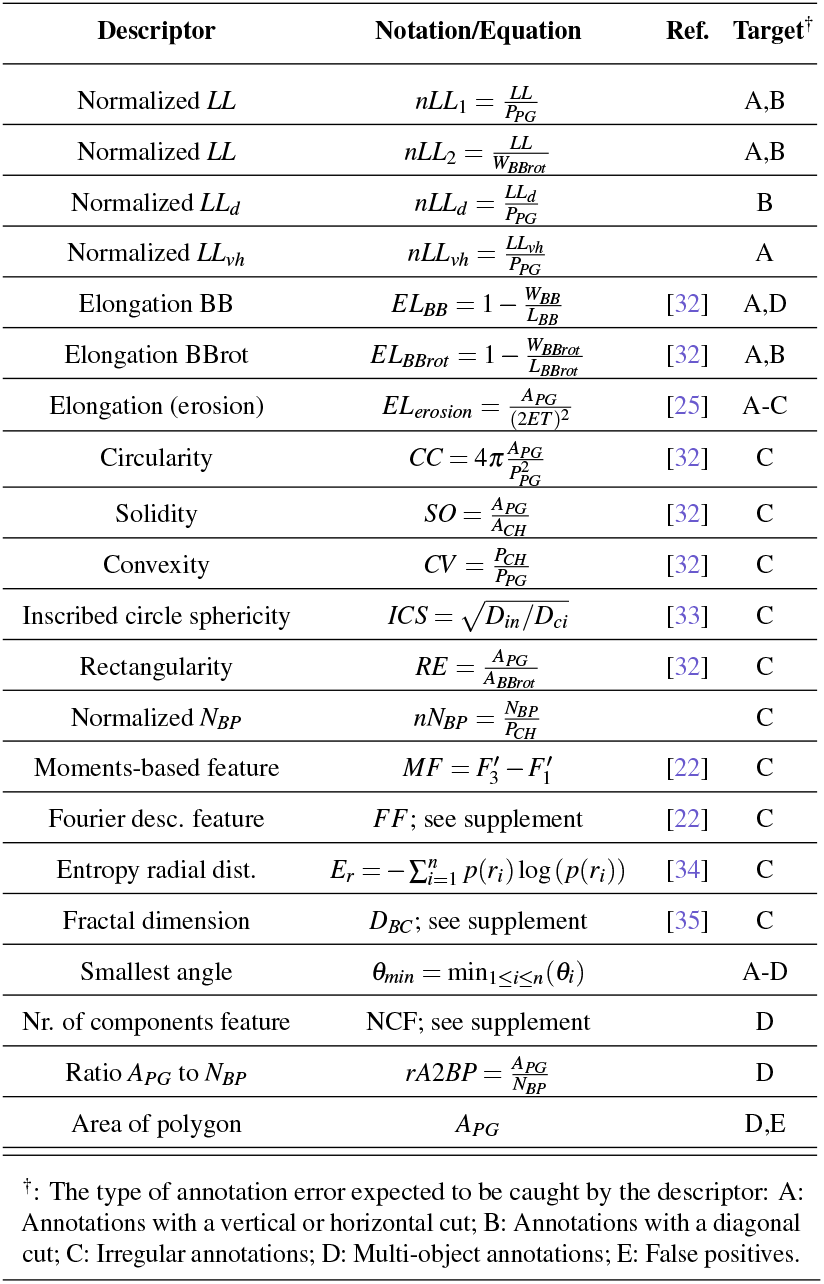
List of shape descriptors.

| Descriptor | Notation/Equation | Ref. | Target<sup>†</sup> |
| --- | --- | --- | --- |
| Normalized $LL$ | $nLL_1 = \frac{LL}{P_{PG}}$ | | A,B |
| Normalized $LL$ | $nLL_2 = \frac{LL}{W_{BBrot}}$ | | A,B |
| Normalized $LL_d$ | $nLL_d = \frac{LL_d}{P_{PG}}$ | | B |
| Normalized $LL_{vh}$ | $nLL_{vh} = \frac{LL_{vh}}{P_{PG}}$ | | A |
| Elongation BB | $EL_{BB} = 1 - \frac{W_{BB}}{L_{BB}}$ | [32] | A,D |
| Elongation BBrot | $EL_{BBrot} = 1 - \frac{W_{BBrot}}{L_{BBrot}}$ | [32] | A,B |
| Elongation (erosion) | $EL_{erosion} = \frac{A_{PG}}{(2ET)^2}$ | [25] | A-C |
| Circularity | $CC = 4\pi \frac{A_{PG}}{P_{PG}^2}$ | [32] | C |
| Solidity | $SO = \frac{A_{PG}}{A_{CH}}$ | [32] | C |
| Convexity | $CV = \frac{P_{CH}}{P_{PG}}$ | [32] | C |
| Inscribed circle sphericity | $ICS = \sqrt{D_{in}/D_{ci}}$ | [33] | C |
| Rectangularity | $RE = \frac{A_{PG}}{A_{BBrot}}$ | [32] | C |
| Normalized $N_{BP}$ | $nN_{BP} = \frac{N_{BP}}{P_{CH}}$ | | C |
| Moments-based feature | $MF = F'_3 - F'_1$ | [22] | C |
| Fourier desc. feature | $FF$; see supplement | [22] | C |
| Entropy radial dist. | $E_r = -\sum_{i=1}^n p(r_i) \log(p(r_i))$ | [34] | C |
| Fractal dimension | $D_{BC}$; see supplement | [35] | C |
| Smallest angle | $\theta_{min} = \min_{1 \leq i \leq n}(\theta_i)$ | | A-D |
| Nr. of components feature | $NCF$; see supplement | | D |
| Ratio $A_{PG}$ to $N_{BP}$ | $rA2BP = \frac{A_{PG}}{N_{BP}}$ | | D |
| Area of polygon | $A_{PG}$ | | D,E |

<sup>†</sup>: The type of annotation error expected to be caught by the descriptor: A: Annotations with a vertical or horizontal cut; B: Annotations with a diagonal cut; C: Irregular annotations; D: Multi-object annotations; E: False positives.

Specifically, to detect incomplete annotations with a clear cut, the project implemented shape descriptors based on the elongation of the (rotated) bounding box (*EL*_*BB*_, *EL*_*BBrot*_), or based on the length of the longest boundary segments (*nLL*_1_, *nLL*_2_, *nLL*_*d*_, *nLL*_*vh*_). To prioritize incomplete annotations with an irregular boundary, the project utilized descriptors for determining shape and boundary roughness, including circularity (*CC*), solidity (*SO*), convexity (*CV*), rectangularity (*RE*), inscribed circle sphericity (*ICS*, Supp. fig. 4), the erosion-based elongation (*EL*_*erosion*_), the entropy of radial distances (*E*_*r*_), a moments-based feature (*MF*), the fractal dimension based on box-counting (*D*_*BC*_, Supp. fig. 5), a Fourier descriptor-based feature (*FF*, Supp. fig. 6), the normalized number of boundary points (*nN*_*BP*_), and the smallest angle between adjacent boundary line segments (*θ*_*min*_). To detect multi-object segmentations, the project employed the raw polygon area (*A*_*PG*_), a measure based on the ratio between area and boundary points (*rA*2*BP*), and an additional novel shape descriptor was developed based on the number of components observed when subjecting the annotation to a distance transform (*NCF*, Supp. fig. 7).

After application to all glomerular segmentations, the values of the shape descriptors were min-max normalized and the *nLL*_1_, *nLL*_2_, *nLL*_*d*_, *nLL*_*vh*_, *EL*_*BB*_, *EL*_*BBrot*_, *EL*_*erosion*_, *nN*_*BP*_, *MF, E*_*r*_, *D*_*BC*_, *NCF, rA*2*BP*, and *A*_*PG*_ metrics were adjusted by subtraction from 1, so that all descriptors would pick up segmentation errors at the leading edge of descriptor values. To detect potential redundancies between shape descriptors, the project conducted a pairwise comparison of descriptor values across all annotations via the `corrplot` library in R (Supp. fig. 8).

### 3.6 Qualitative evaluation of shape descriptors

The performance of the descriptors was initially evaluated in a visual fashion, i.e. (i) comparing their values between normal annotations and annotations from the various error categories, (ii) inspecting the ordering of annotations by individual descriptors, and (iii) evaluating the separation of annotation categories after dimensionality reduction of the feature space produced by the descriptors.

### 3.7 Quantitative evaluation of shape descriptors

Towards a quantitative evaluation of the descriptors, the project then obtained manual labels for a random subset of all segmented objects. Specifically, human annotators were asked to classify each segmentation object as normal or faulty (cut, irregular, and/or multi-object), thus producing a ground truth for each of the selected objects. Utilizing this labeled dataset, an enrichment analysis [36, 37] was then conducted to measure how well the descriptors enriched for these segmentation errors (Supp. fig. 9).

## 4 Results

### 4.1 Visual identification of annotation errors

Based on a visual inspection of the segmentation results, the Histo-Cloud tool [4] demonstrated an excellent performance in detecting and segmenting glomeruli, with the vast majority of annotations appearing virtually error-free. However, given the large number of segmented objects, a small fraction of annotations was found to harbor mistakes and/or inconsistencies. In total, five types of erroneous annotations were identified. Specifically, the first group comprised annotations that were possibly produced by an inconsistent segmentation between adjacent tiling windows, i.e. producing annotations with a clear vertical or horizontal cut (Fig. 2A-B, Supp. fig. 2A-B). In rare cases, annotations also displayed similar cuts, which however were always close to but neither perfectly horizontal nor vertical (Fig. 2C-D, Supp. fig. 2C), but might likely have been caused by a similar issue. Another large fraction of the incomplete annotations displayed more irregular contours (Fig. 2E-F, Supp. fig. 2D), possibly caused by difficulties in detecting glomerular boundaries. The fourth category of errors encompassed annotations in which multiple adjacent glomeruli were segmented as a single object (Fig. 2G-H, Supp. fig. 2E). The fifth group of errors comprised FP detections (Fig. 2I-J). Detailed manual human labeling of a small random subset of objects from each of the five datasets suggested that, on average, around 1-2% of annotations exhibited cuts, around 4-6% were irregular, around 1% were multi-objects, and less than 1% were FP (Supp. fig. 3A).

Finally, an inspection at the WSI-level also suggested the presence of FN predictions (Supp. fig. 3B-D). Specifically, manual human labeling in a subset of WSIs indicated that the percentage of FNs fell between 0% and 15% in most cases (Supp. fig. 3B), and they were mostly accounted for by glomeruli with global sclerosis and/or reduced structural integrity (Supp. fig. 3C-D).

### 4.2 Qualitative evaluation of shape descriptors

To detect the cut, irregular, and multi-object errors, a panel of 21 shape descriptors was established (Table 2). Based on the definition of the individual descriptors, it was expected that some of them might produce strong pair-wise correlations and potential redundancies. However, no identical rankings of annotations were observed (Supp. fig. 8), indicating that all descriptors could be included in the evaluations.

When applied to a representative example selection of correct and faulty annotations, the chosen descriptors were found to produce the expected results in prioritizing individual types of errors (Fig. 4A). Specifically, as compared to a normal annotation, all descriptors displayed lower values in either a cut, irregular, and/or multi-object annotation. These trends were confirmed by visually inspecting annotations sorted based on the values of the individual descriptors (Fig. 4B-D, Supp. fig. 10-14).

**Fig. 4.**
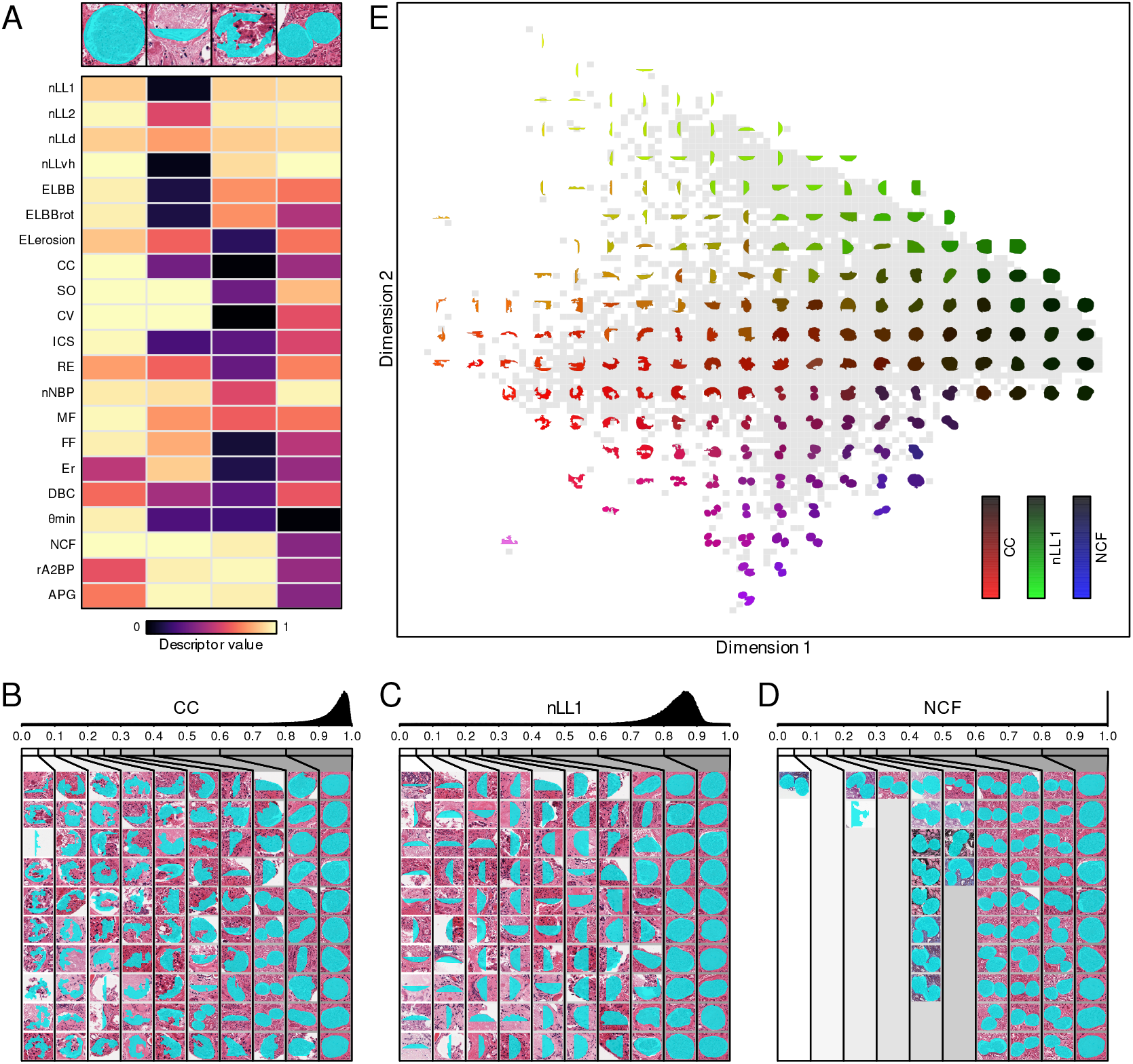
Qualitative evaluation of descriptors. **A)** Heatmap showing the descriptor values between normal annotations and cut/irregular/multi-object annotations. **B-D)** Sorting of annotations by the *CC* (B), *nLL*_1_ (C), and *NCF* (D) descriptors, respectively. The upper panel displays the histogram of values for the respective shape descriptor, while the lower panel displays example annotations across various intervals of the histogram range. **E)** Scatterplot depicting the results of a multi-dimensional scaling down to two components, overlaid with the shapes of example annotations from the various regions of the plot. Shapes were colored according to the values of three descriptors (*CC*: red channel intensity, *nLL*_1_: green channel intensity, and *NCF*: blue channel intensity).

Furthermore, subjecting the *m* × *n*-dimensional data, obtained by computing the *n* = 21 descriptors for all *m* = 168731 annotations, to a principal component analysis (PCA, Supp. fig. 15A-D) [18] or a multi-dimensional scaling (MDS, Fig. 4E, Supp. fig. 15E-F), a good spatial separation could be achieved between normal annotations and each of the three (cut, irregular, multi-object) annotation errors. Investigating the loading of the various descriptors in the PCA plot (Supp. fig. 15A-B), it was further possible to identify the features most relevant in achieving the underlying separation, including e.g. *CC, nLL*_1_, and *NCF* (Fig. 4B-D). Finally, performing a density analysis (Supp. fig. 15E) on the results of the two-dimensional MDS, it was possible to obtain a rough limit on the percentage of annotations possibly exhibiting extensive inconsistencies or faults, which was estimated to be somewhere around 6-8% of all valid annotations (Supp. fig. 15F), consistent with the results from human manual labeling (Supp. fig. 3A).

### 4.3 Quantitative evaluation of shape descriptors

To approach a more exact evaluation of the use of shape descriptors in highlighting segmentation faults, the project then established four datasets of glomerular annotations with an accompanying ground truth, i.e. human-established labels indicating whether the annotation shape was considered normal or faulty (cut, irregular, and/or multi-object; Supp. fig. 3A), where each dataset comprised a unique subset of segmentations across all repositories and stains labeled by one out of four human annotators. Using the labeled datasets, an enrichment analysis [36, 37] was then conducted, evaluating how well segmentation faults were enriched on the leading-edge of a ranking achieved by a descriptor (Supp. fig. 9). The experiment was first conducted separately for each combination of one of the 21 shape descriptors and one of the three error categories (cut, irregular, multi-object) (Supp. fig. 16). A visual inspection suggested that *θ*_*min*_ might perform generally best for prioritizing cut annotations, while *CV* and *NCF* were among the best descriptors for prioritizing irregular and multi-object annotations, respectively (Supp. fig. 16). Specifically, the use of these descriptors resulted in an accumulation of almost all errors in the leading 30% (Cut), leading 20% (Irregular), or leading 5% (Multi-Object) of all annotations (Supp. fig. 17A-C), with very few outliers potentially caused by mislabeling or labeling of negligible inconsistencies by the human annotators (Supp. fig. 17D-F). In order to prioritize any segmentation fault regardless of the error type, the project then combined the three descriptors when sorting annotations (Fig. 5A), again producing an excellent enrichment. Specifically, apart from seven outliers, all errors were again restricted to the leading 30% of the sorted annotations (Fig. 5A). In summary, accounting for a few outliers possibly caused by mislabeling or labeling of minor segmentation inconsistencies (Fig. 5B), the combined ranking with the three selected shape descriptors was considered highly effective at prioritizing any segmentation faults, and the individual descriptors were even more effective at prioritizing irregular and multi-object annotations.

**Fig. 5.**
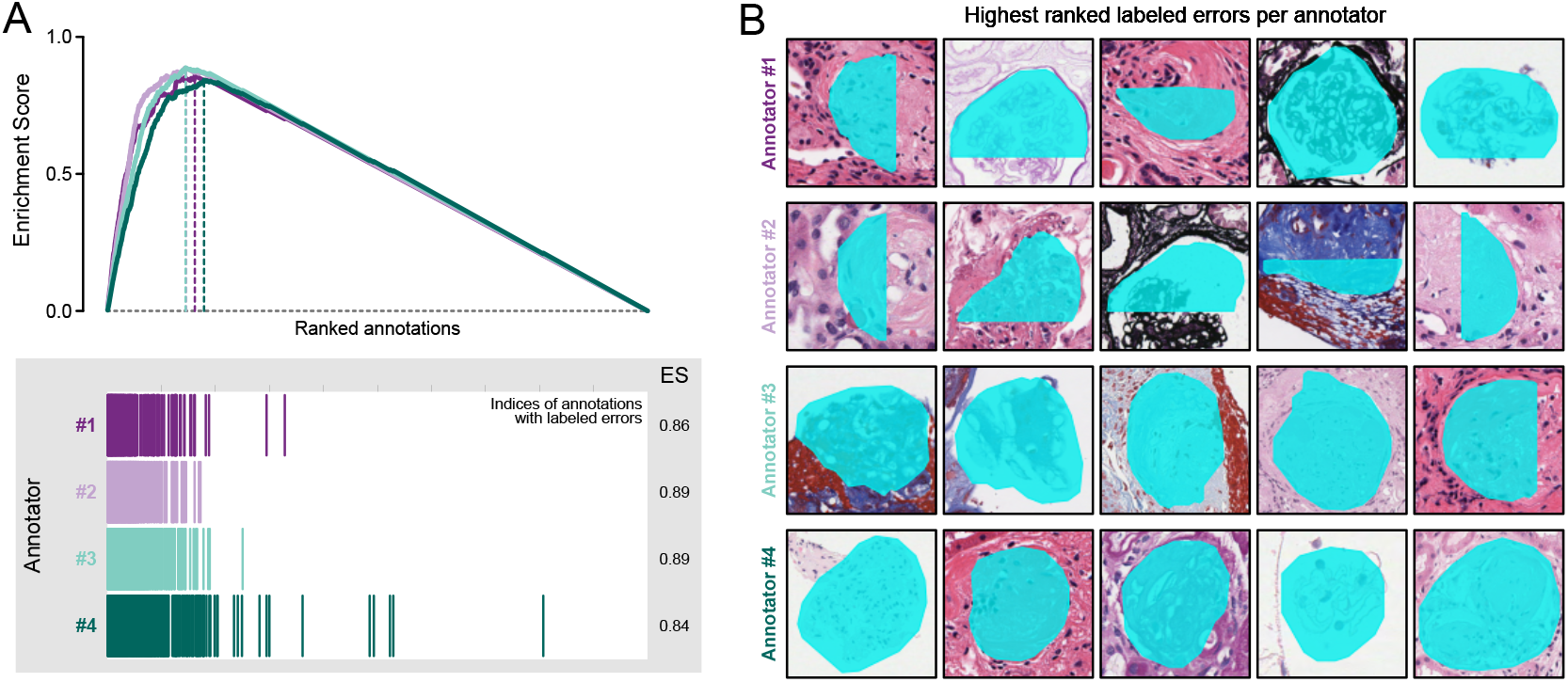
Quantitative evaluation of descriptors. **A)** Enrichment analysis of segmentation faults after sorting annotations using a combination of the *CV, θ*_*min*_, and *NCF* descriptors. Specifically, the 4000 annotations labeled by each annotator were ranked for each of the three descriptors, and then the minimum rank across the three descriptors was used to sort annotations. The bottom panel displays the indices of labeled faults among the sorted annotations, and the upper panel displays the resulting enrichment curves. **B)** The five highest ranked annotations labeled as faulty by each annotator, respectively.

## 5 Discussion

DL-based segmentation represents a crucial step in the automatic characterization of glomerular lesions in nephropathology [13, 38]. However, such segmentations sometimes result in inconsistencies [3, 8, 9] that might require correction before using the output in downstream applications. Manually validating all segmentations would be a rather time-consuming tasks, scaling linearly with the number of objects to be evaluated. Accordingly, recent research has focused on methods for automatically prioritizing [8] and/or automatically correcting potential inconsistencies [8, 9]. While these studies have utilized shape analysis as the core method for the detection of inconsistencies, they only utilized a single shape descriptor/conversion and did not fully investigate which types of segmentation faults can be expected and how such errors would be picked up by the chosen shape descriptor.

To overcome this gap in the literature, the current study conducted a more extensive investigation into the predominant types of faults occurring during glomerular segmentation and investigated a broad panel of shape descriptors in their use to prioritize such mistakes. Specifically, by screening more than 168000 segmentation annotations, the study identified three broad categories of inconsistencies that might be captured through shape analysis. Subsequently, investigating 21 shape descriptors, the study demonstrated that shape analysis is able to produce an estimate on the prevalence of such inconsistencies in a dataset and to prioritize them so that downstream corrections can be focused on the segmentations most demanding of manual validation/correction. Altogether these findings indicate that shape analysis can provide substantial aid in the postprocessing of glomerular segmentations, by estimating error rates and highlighting segmentations that might require manual corrections. Furthermore, the study identified a set of shape descriptors that appear most suitable for such analyses. Among these metrics is a novel descriptor (NCF, Supp. fig. 7), developed during the study, that was shown highly specific for prioritizing annotations harboring multiple adjacent glomeruli segmented as a single object.

Notably, the findings documented in this study are also subject to some limitations. For instance, only a single model was employed for generating glomerular segmentations. Since the start of the project, multiple alternative glomerular segmentation models have been published [5– 7, 10, 11], and it would be relevant to include some of them into a more comprehensive evaluation of glomerular segmentation errors. Nevertheless, based on other publications, it appears evident that the error categories discussed in the current study, including irregular [3, 5, 9], multi-object [8, 10], and cut [3, 5] annotations already encompass the most common faults across segmentation tools. Also, it remains unclear how much the postprocessing of segmentations might affect the descriptors, e.g. when the segmentation boundaries are not smoothened the *θ*_*min*_ descriptor might become less useful.

Importantly, as became apparent in the evaluation of expert-labeled annotations, establishing a reliable ground truth for segmentation errors is difficult and subject to inter-observer variability, which is a common problem in digital pathology. Specifically, the extent of inconsistencies varied substantially across the observed segmentation faults, ranging from minor deviations in a single boundary segment to major mistakes affecting most of the annotation. Considering this gradient, it seemed impossible to establish a universal cut-off between what should be considered an obvious mistake and what would constitute an acceptable annotation. Thus, while there occurred only few obvious and potentially accidental mislabelings of errors (e.g. between error categories or between normal and faulty annotations), the resulting labels were still expected to exhibit intra- and inter-observer variability. The establishment of a segmentation ground truth, rather than the labeling of glomeruli into error categories, might have enabled a more reliable means of measuring the actual accuracy of any segmentation annotation. However, measures such as intersection-over-union or Dice score would not have enabled an actual investigation of the shape descriptors’ abilities to detect certain types of segmentation mistakes.

Furthermore, with respect to the relationship between descriptors and segmentation errors, it should be noted that changes in properties such as circularity, area, or elongation might also be a sign of advancing kidney disease progression [23, 24]. However, based on the analysis of ground-truth labeled glomerular segmentations and of histograms of descriptor values in the current study, we expect that inconsistencies caused by faulty segmentations would typically cause much more pronounced shape deviations than disease-related morphological changes to glomerular shape.

Furthermore, while shape analysis appears highly suitable in identifying some obvious types of segmentation faults, it is clearly not able to address all types of errors, such as false positives and false negatives. Specifically, while a fraction of false positives might also be detected as incomplete/irregular annotations, a truly dedicated detection and correction of false positives and false negatives will likely require a more sophisticated strategy beyond mere shape analysis. For false positives, alternative detection strategies might for instance include color-or texture-based features [39] and/or the use of classification models distinguishing between glomerular and non-glomerular image patches [40, 41]. False negatives in this context would include both (i) annotations that are perfectly regular but too small to capture the entire glomerulus and (ii) glomeruli that are missed entirely. A possible detection strategy for such faults could involve the combination of multiple segmentation models, as e.g. suggested by Yue et al. [6]. For instance, glomeruli found by only a subset of the models might resemble false positives or false negatives and would require manual inspection. For other detections, information from all segmentations might be used to determine the final annotation, after which shape analysis can be utilized to identify any remaining annotation error and initiate downstream manual or automatic corrections. Such a strategy might also be employed using a single model but employing different down-scaling factors during segmentation, which might also affect the ability of the model to detect glomeruli.

Finally, the current study focused entirely on the highlighting of potential segmentation errors, but did not discuss any downstream applications for correcting faulty segmentations. A manual correction often involves the interaction with segmentation results via standard image viewers [42, 43]. Fully automated strategies published so far include for instance taking the convex hull of any segmentation [9] or clustering of segmentations deemed to be multi-objects [8]. Automatic corrections might typically be much faster and reproducible than manual corrections, but also allow little if any control over the correction of individual annotations, thus introducing potential systematic biases. Accordingly, we believe that future approaches will likely benefit from combining automatic and manual corrections. Specifically, combining a voting over multiple segmentation algorithms followed by shape analysis might provide the basis for distinguishing between segmentations that can be effectively validated/corrected in an automatic fashion as compared to predictions that require manual validation by an expert.

## 6 Conclusion

The current study demonstrates that morphometry provides a versatile set of tools for prioritizing certain types of glomerular segmentations faults. In addition to highlighting three common types of segmentation inconsistencies, the study identified three respective shape descriptors that proved highly efficient at detecting these mistakes, enriching virtually all errors in the leading 30% of annotations. Among these metrics was a bespoke shape descriptor designed in the current study specifically to detect multi-object annotations. In summary, the outlined descriptors might be used to highlight annotations in need of manual inspection or correction, thus aiding in (i) the establishment of additional training data for improving the segmentation model or (ii) the curation of glomerular datasets for downstream applications. We believe that the sorting strategy would drastically reduce the time pathologists need to spend on the exhausting validation of segmentation annotations, instead allowing them to focus only on the annotations most likely to contain errors.

## Data availability

Data is available from the corresponding author on reasonable request.

## Author contributions

*Conceptualization*: H.W.; *Data curation*: H.W., J.B., N.M., S.S., S.L.; *Formal analysis*: H.W.; *Funding acquisition*: S.L.; *Investigation*: H.W.; *Methodology*: H.W.; *Project administration*: H.W.; *Resources*: S.L.; *Software*: H.W.; *Supervision*: S.L., H.W.; *Validation*: H.W.; *Visualization*: H.W.; *Writing – original draft*: H.W.; *Writing – review and editing*: H.W., J.B., N.M., S.S., S.L.

## Acknowledgments

The results here are in whole or part based upon data generated by the Kidney Precision Medicine Project (KPMP, https://www.kpmp.org; last accessed/downloaded: March 27, 2025) and the Genotype-Tissue Expression (GTEx, https://www.gtexportal.org; last accessed/downloaded: June 22, 2022). The Kidney Precision Medicine Project is supported by the National Institute of Diabetes and Digestive and Kidney Diseases (NIDDK) through the following grants: U01DK133081, U01DK133091, U01DK133092, U01DK133093, U01DK133095, U01DK133097, U01DK114866, U01DK114908, U01DK133090, U01DK133113, U01DK133766, U01DK133768, U01DK114907, U01DK114920, U01DK114923, U01DK114933, U24DK114886, UH3DK114926, UH3DK114861, UH3DK114915, and UH3DK114937. The Genotype-Tissue Expression Project was supported by the Common Fund of the Office of the Director of the National Institutes of Health, and by NCI, NHGRI, NHLBI, NIDA, NIMH, and NINDS.

