## Supplemental Material for "Morphometry-based detection of deep learning faults in glomerular segmentation"

---

**References** **24**

### 1. Supplementary methods

#### 1.1 Data preparation

##### 1.1.1 Glomerular segmentation

Glomerular candidates in the gathered collection of WSIs were automatically detected and segmented utilizing the Histo-Cloud tool [1], using the *model-Glomeruli-11-13-20* model. KPMP WSIs were processed using default settings, i.e. using a downsampling factor of 2, while the GTEx WSIs were processed using a downsampling factor of 1. Thus, with the original KPMP images coming at 40x magnification and GTEx images coming at 20x magnification, both would be segmented at 20x.

For each WSI, the Histo-Cloud would provide instance segmentation results as a list of annotations, one per detected object and each in the form of a polygon defined by a set of border points. These lists of annotations were merged into a single table across all WSIs, keeping information about the source WSI and adding a global identifier to each detected object.

Importantly, glomeruli were automatically segmented from WSIs without any constraints/selections about regions of interests. Consequently, since WSIs often included more than one tissue section and/or a single biopsy was analyzed using different stains, the resulting data set often contained multiple images (originating from different sections/stains) for the same glomeruli. However, since none of the images were perfectly identical, none of the duplicates were removed.

##### 1.1.2 Generation of image patches

In order to visually process individual segmentations, the project produced markup images, i.e. a separate image patch for each segmented object overlaid with the respective segmentation annotation. Specifically, utilizing the OpenSlide [2] library in Python to read the .svs WSI files, each annotation was used to extract a region of the image centered around the center $(x_c, y_c)$ of the respective polygon, defined as

$$(x_c, y_c) = \left( \frac{min(x_{PG}) + max(x_{PG})}{2}, \frac{min(y_{PG}) + max(y_{PG})}{2} \right),$$

where $x_{PG}$ and $y_{PG}$ are the x and y coordinates of the polygon boundary points. The size of the image patches was defined symmetrically, so that

$$w_{patch} = h_{patch} = \max(w_{PG}, h_{PG}) + 2d,$$

where $w_{patch}$ and $h_{patch}$ are the width and height of the final patch, $w_{PG}$ and $h_{PG}$ are the width and height of a bounding box (with edges parallel to the x and y axes) around the polygon, and $d$ denotes the padding, i.e. the number of pixels that are added on each side of the polygon. The padding was typically chosen to be $d = 16$, $d = 32$, or $d = 64$ pixels for different visualizations or computations. Depending on downstream use, image patches were also downscaled differently before being saved to *.png* files.

#### 1.2 Shape analysis

##### 1.2.1 Detection of invalid polygons

Starting with the raw boundary point coordinates obtained from Histo-Cloud, each annotation was first processed via the *shapely* [3] library in Python, modeling the annotation as a polygon using the *Polygon* function from the *shapely.geometry* module. Subsequently, utilizing the *is_valid* attribute and the *explain_validity* function from the *shapely.validation* module, it was determined if and why a polygon was considered invalid (Supp. Fig. 1). Specifically, among the obtained annotations, there were only two types of invalidities reported by *shapely*: "Self-intersection", i.e. intersecting boundary lines, and "Ring Self-intersections", i.e. intersection at boundary points (Supp. Fig. 1A-D). A subset of these invalidities also included cases in which two adjacent boundary lines displayed an angle of 0°, i.e. indicating that the same line segment is used twice (Supp. Fig. 1B,E). Supplementary figure 1F displays various examples of such invalid polygons across the different data sets included in the study. Importantly, solely using the validity criterion, without using any other shape analyses, all these annotations can be flagged as requiring further inspection and correction. Accordingly, they were excluded from any downstream analyses, which instead focused on the use of shape analysis to detect faulty segmentation annotations (represented by valid polygons).

##### 1.2.2 Conversion into alternative annotation shapes

Having represented each annotation as a polygon through *shapely*, three alternative representations, here respectively referred to as convex hull (CH), bounding box (BB), and rotated bounding box (BBrot), were obtained through the *shapely* attributes *convex_hull*, *envelope*, and *minimum_rotated_rectangle*, respectively.

##### 1.2.3 Computation of basic geometric properties

Having obtained the various annotation shape representations as *shapely* objects, several geometric properties could be directly extracted, including the number of boundary points ($N_{BP}$), the perimeter, and the area of an annotation. Specifically, the latter two measures were extracted via the *length* and *area* attributes and multiplied by *res* and $res^2$, respectively, where *res* is the image resolution in $\mu m/pixel$ automatically extracted from the .svs files using the *openslide.mpp-x* property of the *OpenSlide* library in Python. For the rectangular annotations (BB and BBrot), the width and length was taken to be the smaller and larger side length, respectively.

In order to compute radial distances ($r_i$) from each boundary point $p_i = (x_i, y_i)$ to the polygon centroid $(\bar{x}, \bar{y})$, the first step was to acquire the centroid coordinates via the *centroid* attribute of the *shapely* object. Subsequently, distances between boundary points and centroid were calculated using the *distance* function of *shapely*.

The longest line segments (general, horizontal, vertical, diagonal) were determined by simply iterating through each pair of adjacent points, computing the length of the line segment again through the *distance* function, and keeping track of the longest length of segments with the respective orientation.

The angle at a boundary point $p_i$ was obtained by first computing the outside and inside angles between the two adjacent line segments, and then keeping the minimum of those two angles.

##### 1.2.4 Computation of shape descriptors

###### *1.2.4.1 Inscribed circle sphericity*

The inscribed circle sphericity is defined as [4]

$$ICS = \sqrt{\frac{D_{in}}{D_{ci}}} = \sqrt{\frac{2R_{in}}{2R_{ci}}},$$

where $D_{in}$ and $R_{in}$ are the diameter and radius, respectively, of the maximum inscribed circle, and $D_{ci}$ and $R_{ci}$ are the diameter and radius, respectively, of the minimum circumscribed circle for the shape.

$D_{in}$ and $D_{ci}$ were computed through the use of functions available from the cv2 OpenCV [5] library (Supp. Fig. 4). Specifically, each polygon (Supp. Fig. 4A-C) was first converted into a corresponding mask image (Supp. Fig. 4D-F). Subsequently, the *distanceTransform* function was applied to the mask, and the location and value of the maximum result were extracted as the center and radius, respectively, of the maximum inscribed circle (Supp. Fig. 4G-I). The center and radius of the minimum circumscribed circle (Supp. Fig. 4J-L) were instead obtained by applying the *minEnclosingCircle* function on the contours of the mask.

###### *1.2.4.2 Fractal (box counting) dimension*

The box counting dimension, $D_{BC}$, was obtained following the description by Wu et al. [6], based on counting the number $N_{BC}(\varepsilon)$ of grid boxes of size $\varepsilon$ intersected by the polygon boundary of an annotation. Specifically, $D_{BC}$ was computed by counting $N_{BC}(\varepsilon)$ for a sequence of $\varepsilon$ values, and then recording the absolute value of the slope of the line fitted to the curve obtained when plotting $\ln(N_{BC}(\varepsilon))$ as a function of $\ln(\varepsilon)$ [6].

To perform this sampling, the project selected a logarithmic sequence of 20 $\varepsilon$ values in the interval $[2^0, 2^5]$ using the *logspace* function. Subsequently, a matrix of the boundary pixels of the polygon (Supp. fig. 5A-C) was extracted using the *polygon_perimeter* function from the *skimage.draw* library, and $N_{BC}(\varepsilon)$ was computed after rebinning/rescaling the boundary pixel matrix using $\varepsilon$ as downsampling factor (Supp. fig. 5D-L). Finally, the value of $D_{BC}$ was then taken to be the negative value of the slope of the line fitted to the curve between $N_{BC}$ and $\varepsilon$ in log-log space (Supp. fig. 5M-O).

###### *1.2.4.3 Moments-based feature*

Adhering to the definition and documentation provided by Shen et al. [7], a moments-based feature (MF) was computed as follows. Utilizing the radial distances $\mathbf{r} = (r_1, \cdots, r_{n-1}, r_n)$ defined above, the first step was to compute the $1th$ moment, which is simply the mean across the radial distances, i.e. defined as [7]

$$m_1 = \frac{1}{n}\sum_{i=1}^{n} r_i,$$

Subsequently, two normalized features, $F_1'$ and $F_3'$, were computed as [7]

$$F_1' = \frac{\left(\frac{1}{n}\sum_{i=1}^{n}(r_i - m_1)^2\right)^{1/2}}{m_1},$$

$$F_3' = \frac{\left(\frac{1}{n}\sum_{i=1}^{n}(r_i - m_1)^4\right)^{1/4}}{m_1},$$

and the final Moments-based feature was then obtained as [7]

$$MF = F_3' - F_1'.$$

###### 1.2.4.4 *Fourier descriptor feature*

Similarly to the Moments-based feature, the computation of the Fourier descriptor feature (FF) was also solely based on the definition by Shen et al. [7]. Specifically, as suggested by the authors, the first step was to ensure that the number of boundary points ($n$) was a power of 2, which was achieved by resampling boundary points after interpolating across boundary line segments to achieve a power of 2 at least as large as the original number of boundary points (Supp. Fig. 6A-F). Next, each of the obtained boundary points $p_i = (x_i, y_i)$ was represented by a complex number [7]

$$z_i = x_i + jy_i.$$

Subsequently, the *fft* function from the *np.fft* module in Python was utilized to compute the Fast Fourier Transform of the sequence of complex numbers, yielding $n$ Fourier descriptors, $FD(i)$; $i = 0, 1, \cdots, n-1$ (Supp. fig. 6G-I). The corresponding normalized Fourier descriptors, $nFD(k)$; $k = -n/2+1, \cdots, 0, \cdots, n/2$ (Supp. fig. 6J-L), were then obtained as [7]

$$nFD(k) = \begin{cases} 0, & \text{if } k = 0, \\ FD(k)/FD(1), & \text{if } k > 0, \\ FD(k+n)/FD(1), & \text{if } k < 0. \end{cases}$$

Finally, the Fourier descriptor feature was then computed as [7]

$$FF = \frac{\sum_{k=-n/2+1}^{n/2} |nFD(k)|/|k|}{\sum_{k=-n/2+1}^{n/2} |nFD(k)|}.$$

###### 1.2.4.5 *Number of components based on distance transform*

To obtain a metric based on an estimate of the number of objects in an annotation, the project started by performing a distance transform on each annotation (Supp. Fig. 7), utilizing functions from the cv2 OpenCV library in Python. Specifically, each polygon (Supp. Fig. 7A-C) was first converted to a corresponding mask image (Supp. Fig. 7D-F). Subsequently, the mask was subjected to a distance transformation (*distanceTransform* function; using a mask size of 5), and then nine different thresholds were applied (*threshold* function; using threshold values equal to $x \max_{dist}$, where $x \in [0.1, 0.2, \cdots, 0.8, 0.9]$ and $\max_{dist}$ is the maximum value of the distance transform). For each threshold, the resulting image was first postprocessed with two iterations of an opening operation (*morphologyEx* function; using a $3 \times 3$ kernel of ones), and then the number of separate components was obtained as the number of independent connected components (*connectedComponents* function, Supp. Fig. 7G-L), and the maximum number of components ($NC$) across the nine thresholds was recorded. Finally, as very irregular annotations of a single glomerulus can yield similarly high numbers of components as a multi-object segmentation (Supp. Fig. 7K-L), the final metric (number of components-based feature; $NCF$) for a polygon $i$ was instead defined as

$$NCF(i) = \begin{cases} A_{PG}(i), & \text{if } NC(i) > 1, \\ min_j(A_{PG}(j)), & \text{if } NC(i) = 1, \end{cases}$$

thus giving the lowest value to regular annotations of single glomeruli, and expected to separate single glomeruli with multi-component annotations (low area values) from multi-objects with multi-component annotations (larger area values).

###### 1.2.4.6 *Elongation (erosion-based)*

In addition to elongation measures based on the aspect ratio of the (rotated) bounding box, which fail to e.g. pick up very elongated but curved objects, the study also employed a different measure of elongation based on the area ($A_{PG}$) and the erosion thickness (ET) of the polygon, which measures the number of erosion steps that can be applied to an object before it vanishes [8]. In the current project, ET was estimated again from mask images, by applying erosion operations via the *erosion* function (using a $3 \times 3$ kernel of ones) from the cv2 OpenCV library and recording the number of iterations needed before all mask pixels disappeared. Subsequently, the erosion-based value of elongation was computed as [8]:

$$EL_{erosion} = \frac{A_{PG}}{(2ET)^2}.$$

#### 1.3 Qualitative analysis of shape descriptors

##### 1.3.1 Rank correlation between descriptors

Having implemented a number of descriptors, the next step, before starting with any evaluations of their usability, was to check for putative redundancies. Specifically, the main purpose of the current study was to identify metrics able to sort annotations in a way that would enrich for faulty annotations. Thus, if two descriptors would produce an identical ordering of annotations, one of them would be redundant and could be excluded. To test for a possible redundancy, the descriptors were applied to all annotations and the Spearman rank correlation coefficient was computed between each pair of descriptors and across all annotations (Supp. Fig. 8). As none of the descriptors showed a perfect rank correlation (Supp. Fig. 8), i.e. no apparent redundancies were detected, all descriptors were retained for subsequent analyses.

##### 1.3.2 Dimensionality reduction

Based on an individual inspection of the ordering achieved by the various descriptors, most seemed to work well in enriching for a particular type of error (Fig. 4B-D, Supp. Fig. 10-14). To gain a better insight into their usability for separating between normal annotations and the various types of erroneous annotations, and to potentially select the most relevant metrics, the project then employed two types of dimensionality reduction, i.e. a principal component analysis (PCA, Supp. Fig. 15A-D) and a multi-dimensional scaling (MDS, Fig. 4E, Supp. Fig. 15E-F). Both analyses were conducted in R, using the *prcomp* function for PCA and the *interpolation_mds* function from the *bigmds* library for MDS, respectively.

In the PCA analysis, assuming three major types of errors (irregular annotations, cut annotations, multi-object annotations), particular focus was placed on the three first principal components. Plotting the first against the second (Supp. Fig. 15A,C) or third (Supp. Fig. 15B,D) component, it was evident that the three components were able to clearly separate between the three types of errors, and a study of the underlying biplots (Supp. Fig. 15A-B) also allowed to determine the relevance of the individual descriptors in this overall separation. For instance, descriptors like the $nLL_1$, $nLL_2$, $EL_{BB}$, and $EL_{BBrot}$ appeared to play a major role in the separation of the cut annotations (Supp. Fig. 15A), while the $NCF$ descriptor displayed the strongest alignment with the separation of the multi-object annotations (Supp. Fig. 15B). The variation along the first principal component (irregular annotations) was associated with various descriptors, including e.g. $CC$, $CV$, $SO$, and $RE$ (Supp. Fig. 15A-B).

A similar separation between error types was also seen when performing the MDS down to only two dimensions (Fig. 4E), which further enabled a density analysis (Supp. Fig. 15E) and thus a rough estimation of the percentages of correct versus faulty annotations (Supp. Fig. 15F). Specifically, the densest area of the MDS plot, likely harboring only regular/normal polygons, appeared to include between 92-94% of all annotations, suggesting that only roughly 6-8% of annotations actually presented with any type of obvious fault.

#### 1.4 Quantitative analysis of shape descriptors

##### 1.4.1 Establishing of ground truth labels for annotation faults

To test the ability of individual descriptors to detect segmentation faults, the project required access to a dataset with definitive information about which annotations actually exhibited such faults. For that purpose, four human annotators, one biomedical researcher (Annotator #1) and three nephropathologists (Annotators #2-#4), were provided with predicted glomerular segmentations and were then tasked to establish ground truth labels for these data, i.e. to indicate for each annotation whether it was cut, irregular, and/or multi-object, or neither of those. Each annotator was given a unique, random selection of 4000 annotations, with 800 annotations each from the GTEx, KPMP-HE, KPMP-PAS, KPMP-SIL, and KPMP-TRI datasets.

The labeling of segmentation errors turned out to be not straightforward, as the boundaries between faulty and acceptable annotations or between the various error categories was not always clear. This difficulty was expected, considering that there would be a gradient from a completely regular to a highly irregular annotation. Accounting for these difficulties, annotators were thus asked to label only obvious mistakes, for instance by labeling only inconsistencies that affected a substantial fraction (e.g. >10%) of the glomerular area. To further counteract labeling mistakes, labeling was conducted in two rounds; in the first round, annotators were asked to label all provided annotations, while in the second round, annotators where then tasked to review and either accept or adjust all previously identified errors. After the second round of labeling, one annotator still voiced uncertainty with respect to some annotations. In a followup discussion, these cases were identified as mostly (i) annotations that exhibited so small or so indistinct inconsistencies that it was questionable whether they should be labeled as faulty at all, and (ii) annotations where it was questionable whether they exhibited the cut and irregular or only the irregular pattern. In consensus with the annotator, it was decided to consider the first group of annotations as non-faulty, while annotations in the second group were all categorized as irregular only.

##### 1.4.2 Enrichment analyses

One of the core objectives of the project was to identify shape descriptors able to sort annotations so that errors would exclusively appear on the leading edge of the ordering. To evaluate the ability of descriptors to achieve such an ordering, the project utilized an enrichment analysis directly based on the popular Gene Set Enrichment Analysis (GSEA)[9]. Specifically, the procedure for evaluating the enrichment for a single descriptor was equivalent to what was previously described in Weishaupt et al. [10]:

1. Let $A$ denote the set of all annotations and let $P \subset A$ be the subset of annotations positive for a certain type of error (ground truth labels, known beforehand).
2. Sort all annotations in increasing order based on the values of the respective descriptor, and let $a_1, a_2, \cdots, a_{|A|}$ denote the sequence of annotations as they occur in the ordered list. The bottom panel in supplementary figure 9A illustrates the descriptor values of a set of annotations after sorting, while the middle panel in supplementary figure 9A illustrates the labels (faulty or non-faulty) of the ordered annotations.
3. Starting with an enrichment score $ES(0) = 0$, iterate over all annotations in the ordered list and compute the running sum:

$$ES(i) = \begin{cases} ES(i-1) + \frac{1}{|P|}, & \text{if } a_i \in P, \\ ES(i-1) - \frac{1}{|A|-|P|}, & \text{if } a_i \notin P. \end{cases} \qquad i = 1, 2, \cdots, |A|.$$

4. Record the maximum positive value of the score observed during this process as

$$ES_{max} = \max_j (ES(j)).$$

   Specifically, as all metrics were defined such that they would produce a lower value for a given annotation error, we were explicitly interested only in the enrichment of errors in the annotations with the low metric scores, where the best possible enrichment produces a separation of 1 from the x-axis (Supp. fig. 9B).

The final enrichment score for a shape descriptor was then the value of $ES_{max}$, with a value closer to 1 indicating a better enrichment.

### 2. Supplementary figures

#### Supplementary figure 1

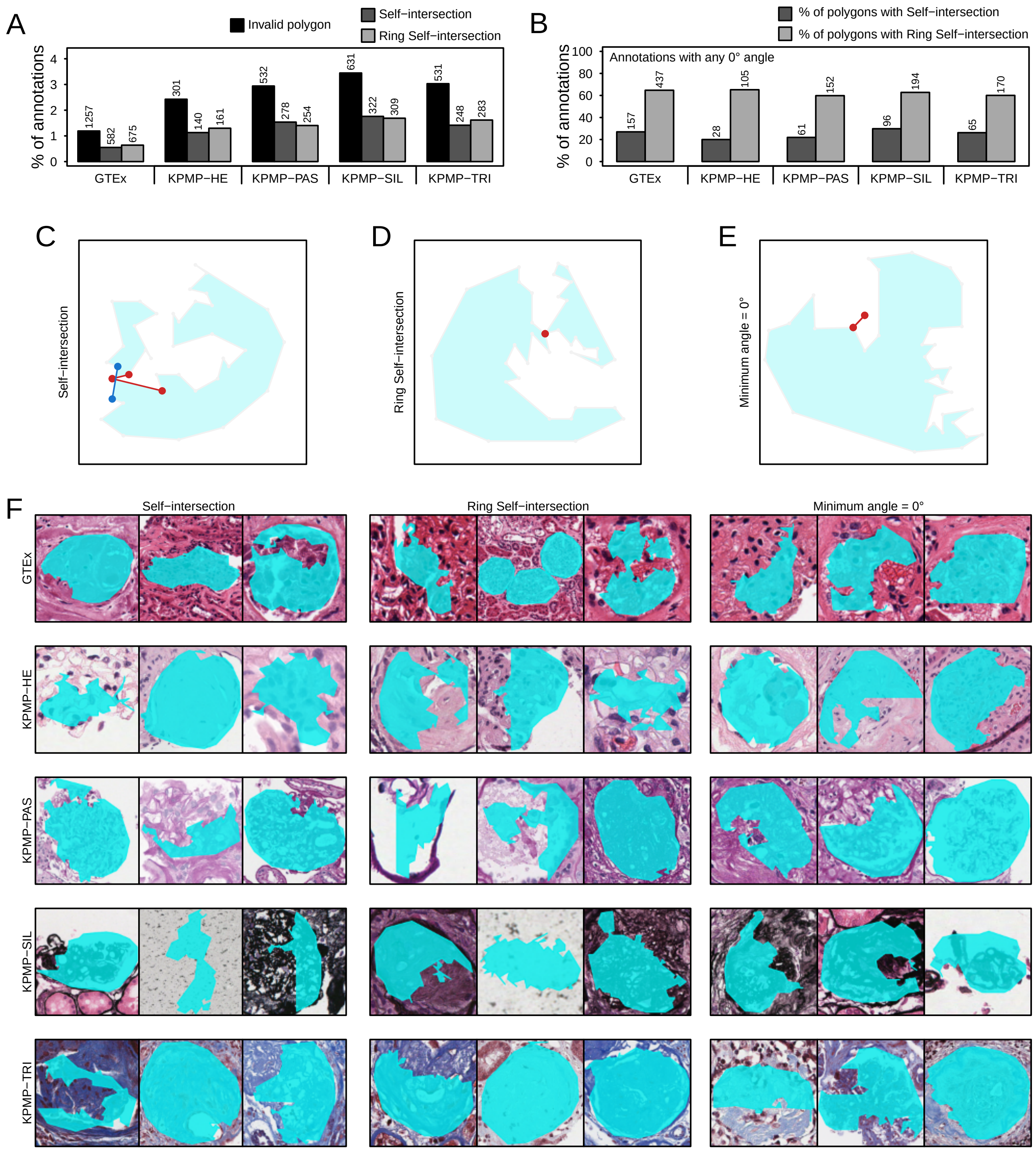

**Supp. Fig. 1.** Investigation of invalid polygons. **A)** The percentages and absolute numbers of annotations represented by invalid polygons per dataset, shown with respect to the two types of invalidity reasons: Self-intersections and ring self-intersections. **B)** Percentages and numbers of annotations with a minimum angle of 0 degrees between any two adjacent boundary line segments among the annotations with self-intersections and ring self-intersections, respectively. **C-E)** Illustrations of a self-intersection invalidity (C), a ring self-intersection invalidity (D), and a 0 degree angle between two adjacent line segments (E). **F)** Examples of annotations exhibiting at least one of the three invalidity types, displayed according to invalidity type (columns) and across the five datasets (rows).

**Supplementary figure 2**

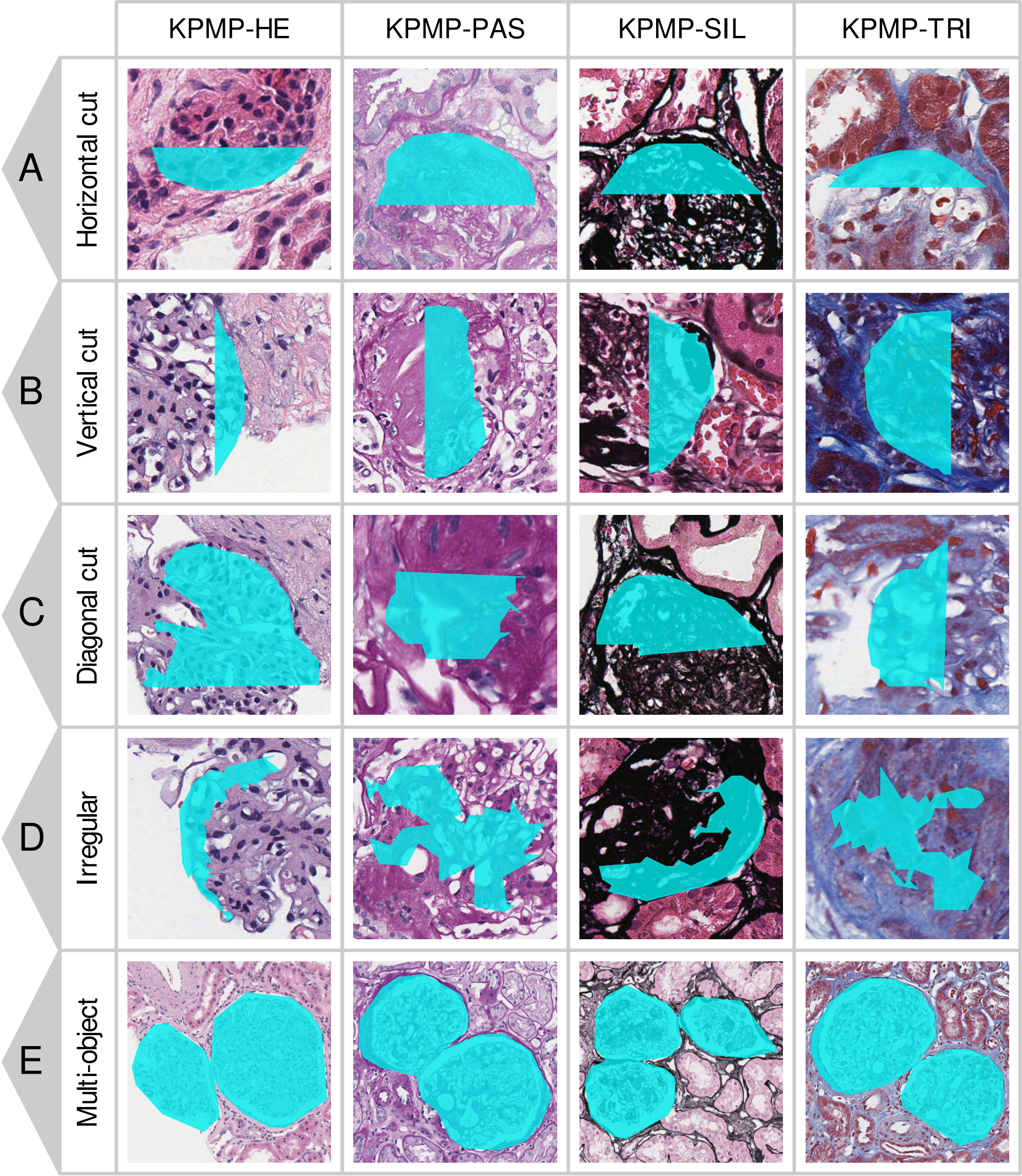

**Supp. Fig. 2.** Examples of faulty segmentation annotations in the KPMP WSIs, including examples of annotations with an abrupt horizontal (A), vertical (B), or diagonal (C) cut, irregular annotations (D), and annotations including multiple objects (E), shown separately for each of the four stains.

#### Supplementary figure 3

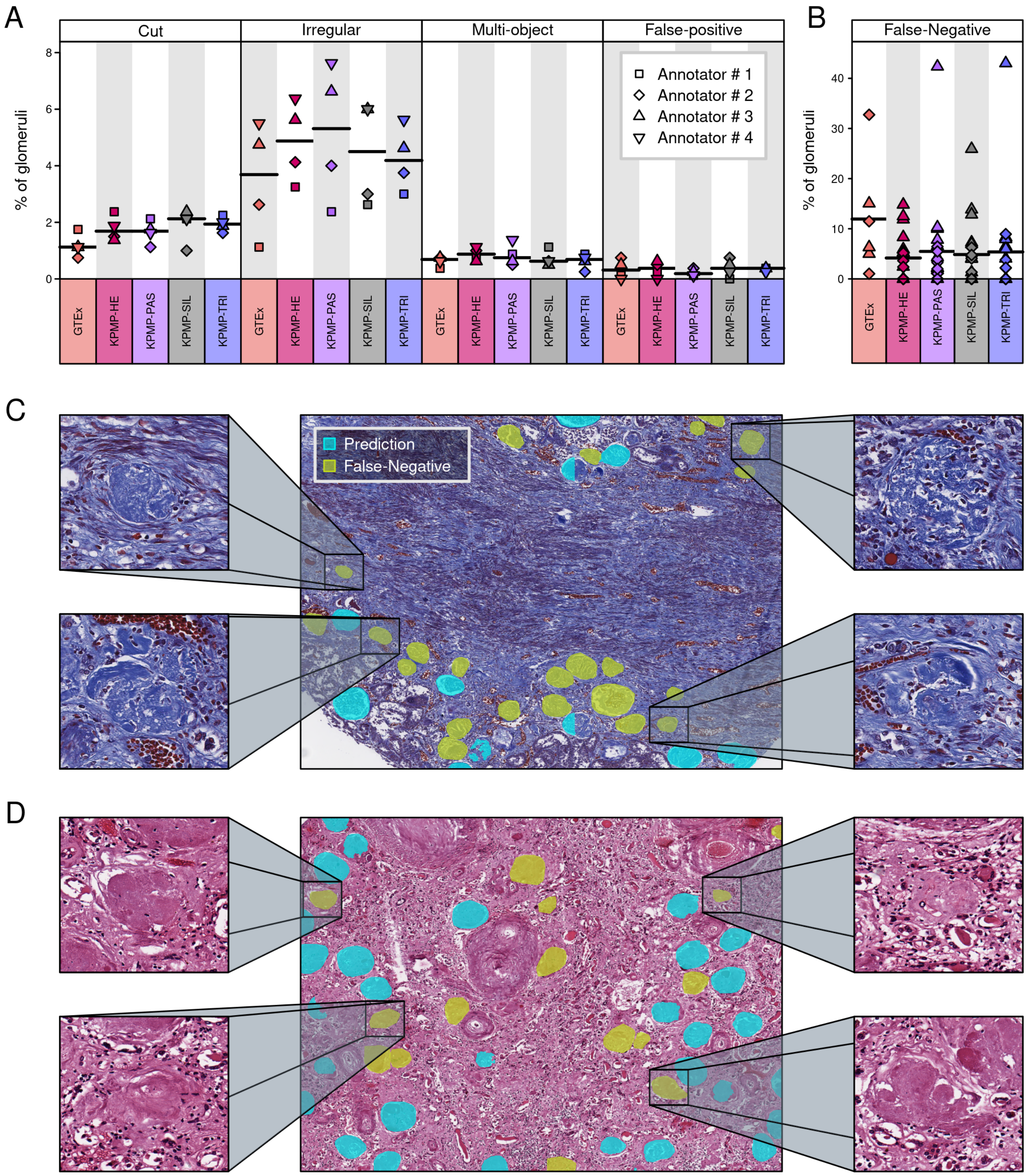

**Supp. Fig. 3.** Quantification of incorrect segmentations. **A)** Percentages of cut, irregular, multi-object, and FP annotations across the different datasets and across the four annotators. Each annotator labeled 800 annotations per dataset. The black horizontal lines indicate the mean values across annotators. **B)** Percentages of FN annotations as estimated by two annotators across 43 WSIs each. **C-D)** Examples of FN glomeruli in a KPMP-TRI (C) and a GTEx (D) WSI.

#### Supplementary figure 4

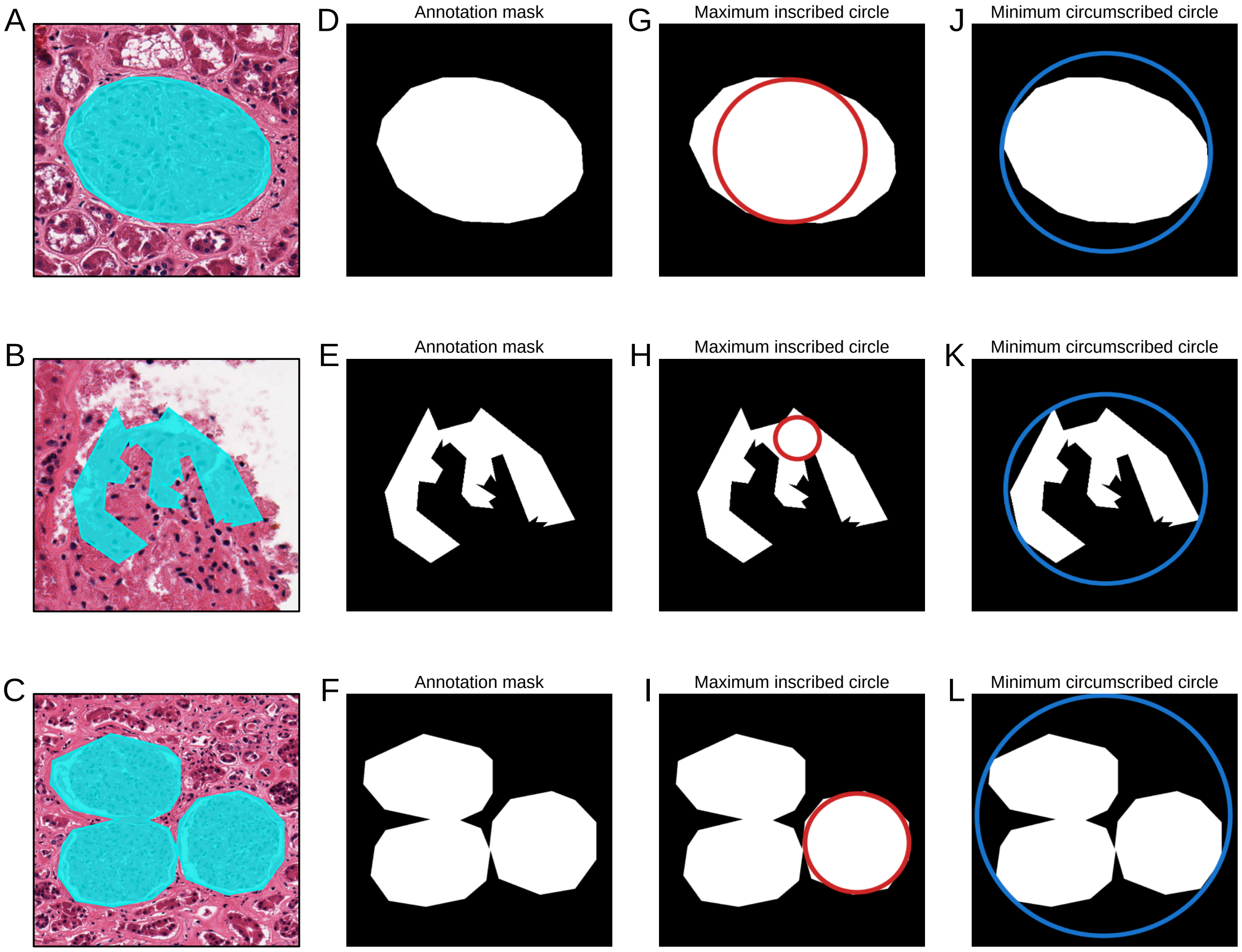

**Supp. Fig. 4.** Obtaining the maximum inscribed and minimum circumscribed circles for annotations. **A-C)** Three different annotations produced by the Histo-Cloud tool. **D-F)** Annotations converted to masks, i.e. white pixels correspond to the area enclosed by the annotation. **G-I)** Maximum inscribed circles with centers and radii obtained as the maximum location and value, respectively, of a distance transformation applied to the mask (*distanceTransform* function followed by *minMaxLoc* function; cv2 OpenCV). **J-L)** Minimum circumscribed circles with centers and radii computed via the *minEnclosingCircle* function from the cv2 OpenCV library.

#### Supplementary figure 5

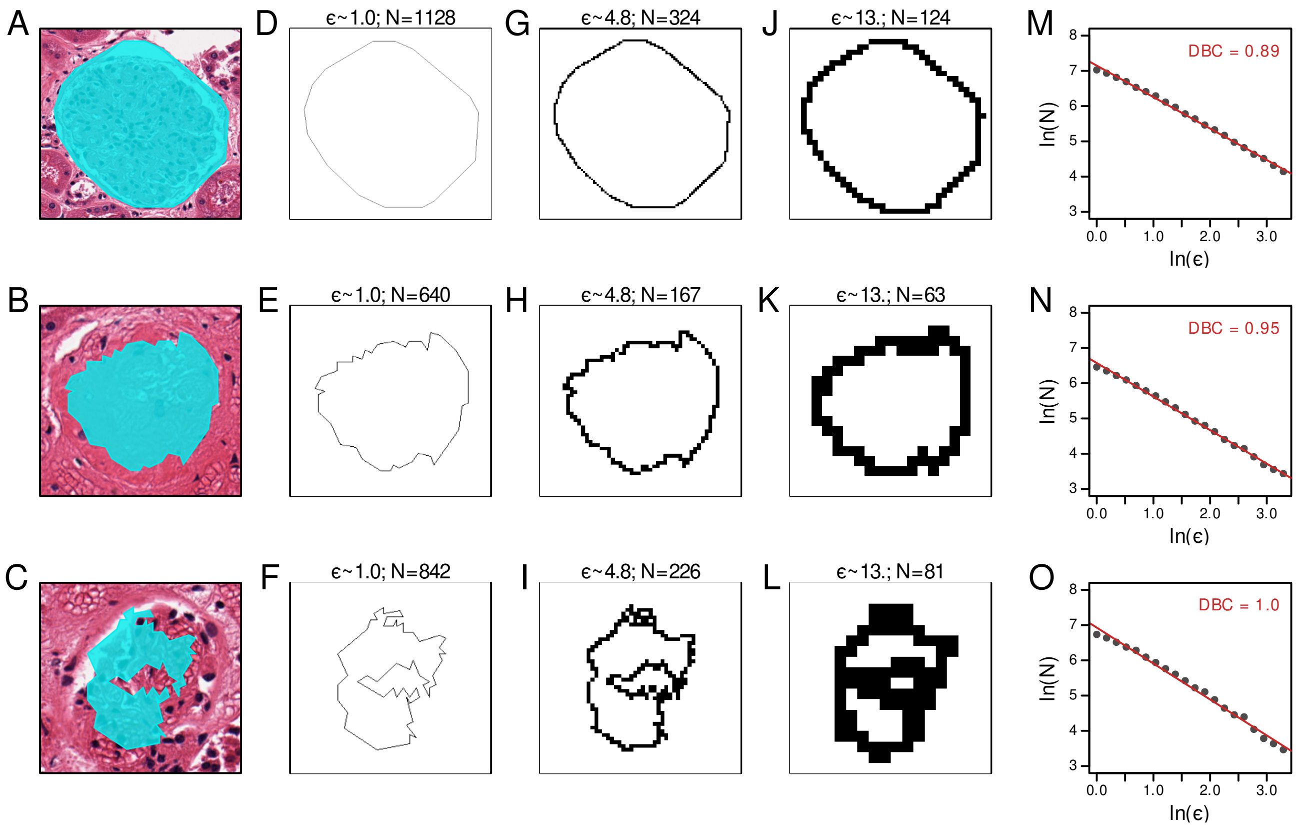

**Supp. Fig. 5.** Fractal (box counting) dimension. **A-C)** Three different annotations reported by Histo-Cloud. **D-F)** The contour of the annotation discretized into $N$ boxes with size $\varepsilon \sim 1.0$ pixels. **G-I)** Discretization into $N$ boxes at $\varepsilon \sim 4.8$ pixels. **J-L)** Discretization into $N$ boxes at $\varepsilon \sim 13$ pixels. **M-O)** The log-log plot showing the relationship between box sizes and number of boxes intersected by the annotation contour.

#### Supplementary figure 6

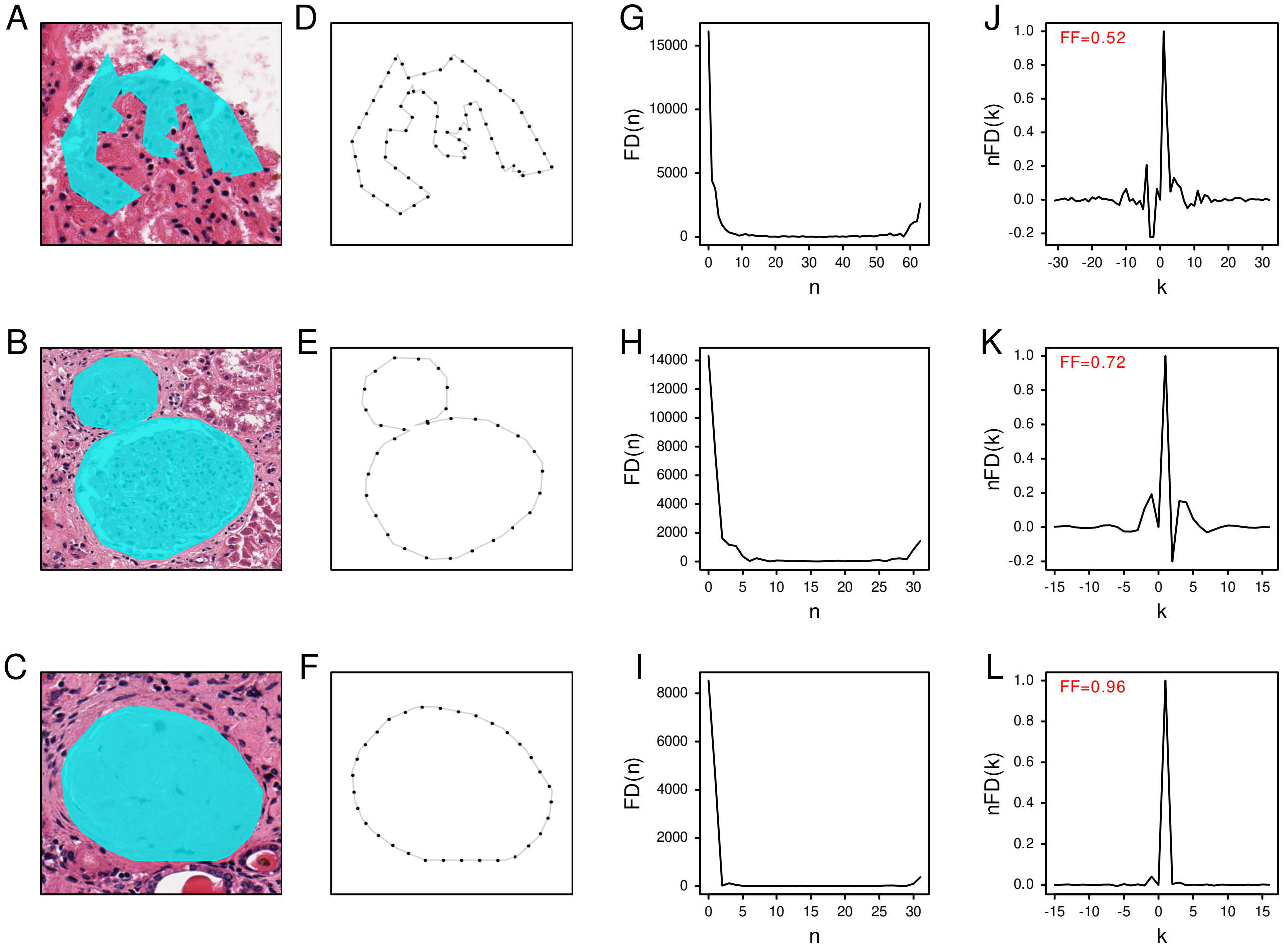

**Supp. Fig. 6.** Fourier descriptor feature computed according to Shen [7]. **A-C)** Three different annotations reported by Histo-Cloud. **D-F)** Interpolation of boundary to get $n$ boundary points (black dots), where $n$ is the smallest power of 2 that is equal to or greater than the original number of boundary points. **G-I)** The raw Fourier descriptors ($FD$) obtained through FFT computation. **J-L)** The normalized Fourier descriptors ($nFD$) plotted against the normalized frequencies ($k$). The resulting value of the Fourier descriptor feature ($FF$) is shown in red.

#### Supplementary figure 7

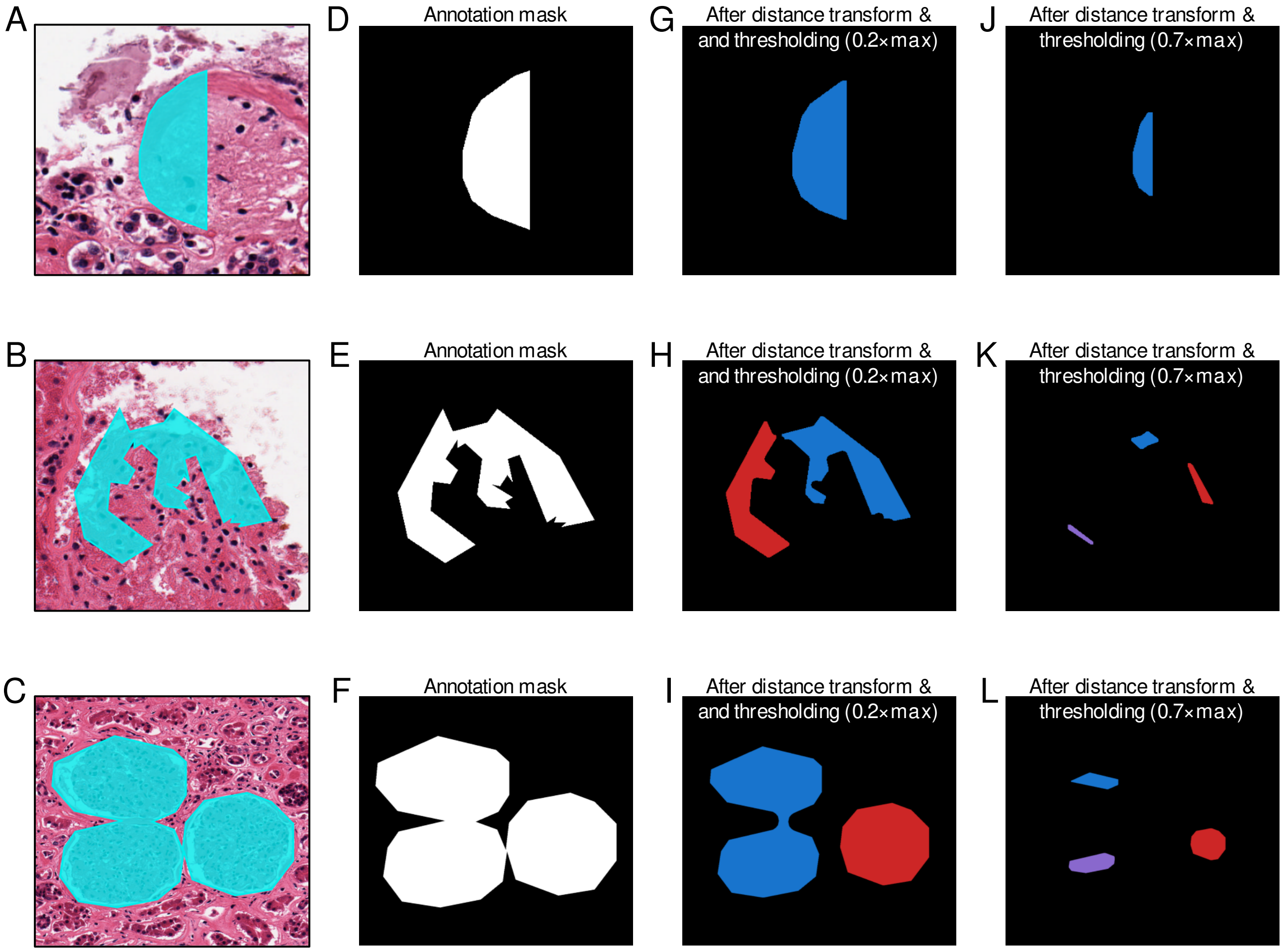

**Supp. Fig. 7.** Distance transform and erosion. **A-C)** Three different annotations reported by Histo-Cloud. **D-F)** Annotations converted to masks, i.e. white pixels correspond to the area enclosed by the annotation. **G-L)** The separate (different colors) components detected after subjecting the mask to a distance transform, a thresholding, using either a threshold equal to 0.2 (G-I) or 0.7 (J-L) times the maximum value of the distance transform, respectively, and a final morphological opening.

#### Supplementary figure 8

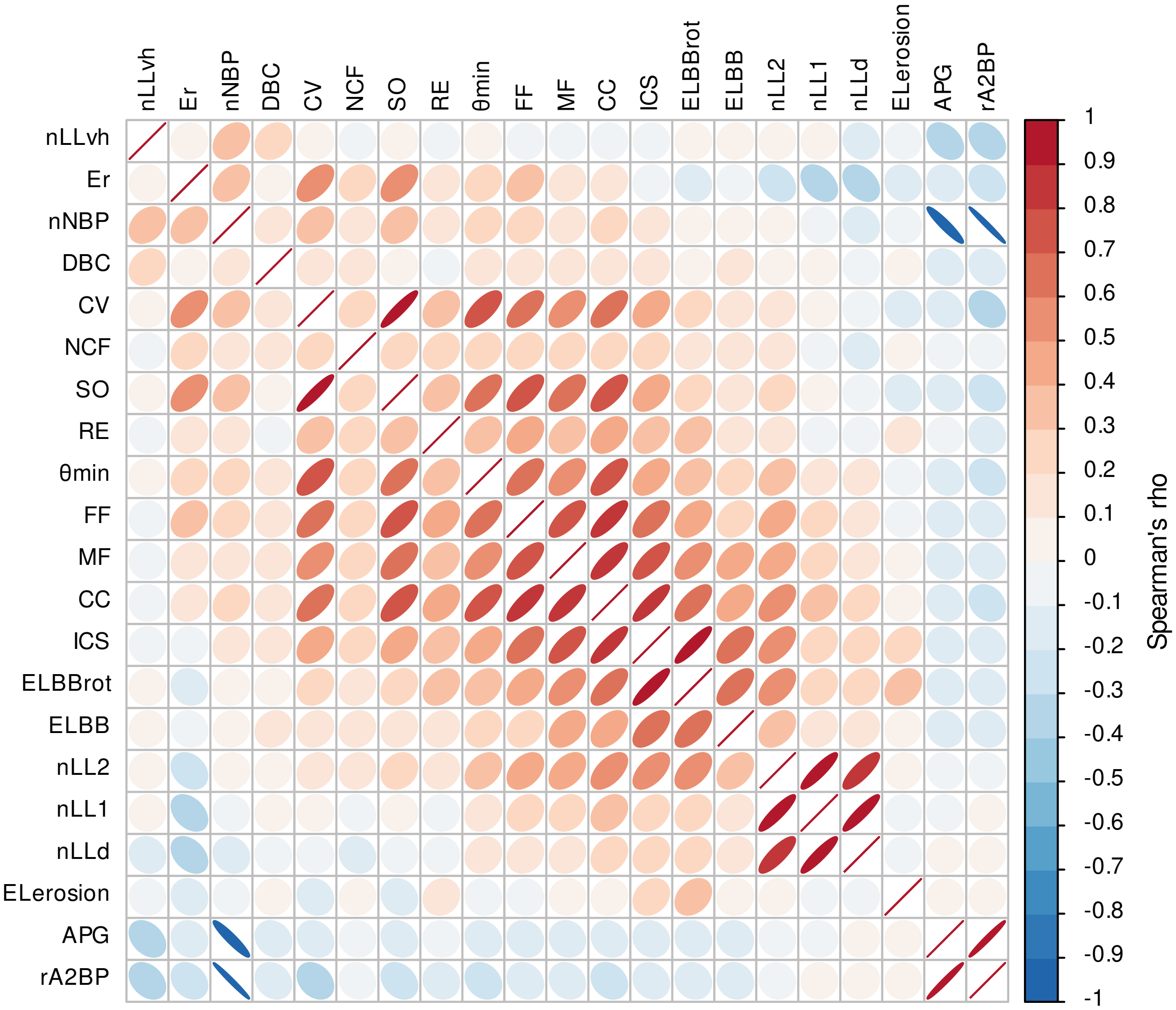

**Supp. Fig. 8.** Correlation matrix, plotted via the *corrplot* library in R, displaying the pairwise Spearman correlation coefficients between shape descriptors across all (n=168731) segmented objects.

#### Supplementary figure 9

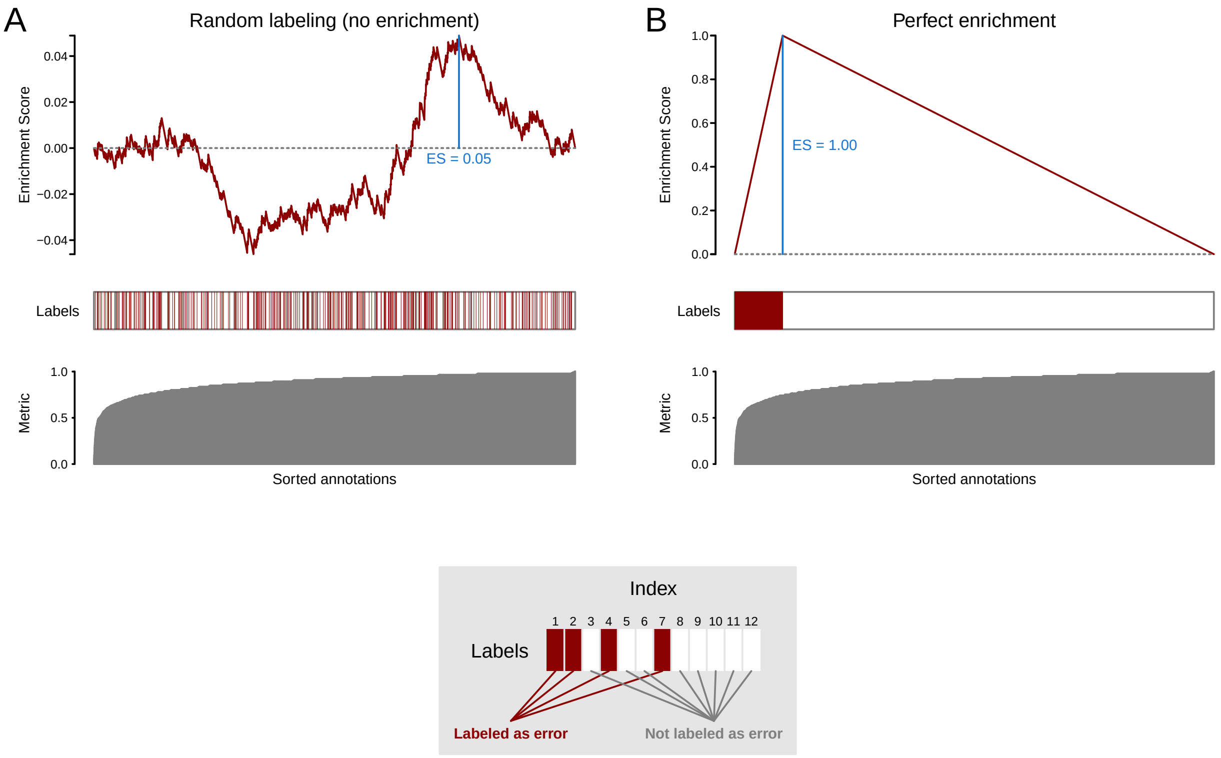

**Supp. Fig. 9.** Illustration of enrichment analysis and generation of enrichment curves. **A)** Enrichment analysis in a scenario where no enrichment is apparent, i.e. annotations are randomly labeled as faulty or non-faulty. **B)** Enrichment analysis in a scenario with a perfect enrichment, i.e. all faulty annotations are on the leading edge. The bottom panel in each subfigure illustrates the value of the shape metric after sorting all annotations based on that descriptor. The middle panel indicates the label (faulty or non-faulty) for each index in the ordered sequence of annotations, where a red line indicates a faulty annotation at the respective index (see also legend for illustration of labeling). The top panel illustrates the resulting enrichment curve (red line), with the final enrichment score (ES) determined as the maximum positive value of the curve (indicated by the vertical blue line). All data in the figure are simulated for the purpose of visualization. See also Subramanian et al. [9] and Weishaupt et al. [10].

#### Supplementary figure 10

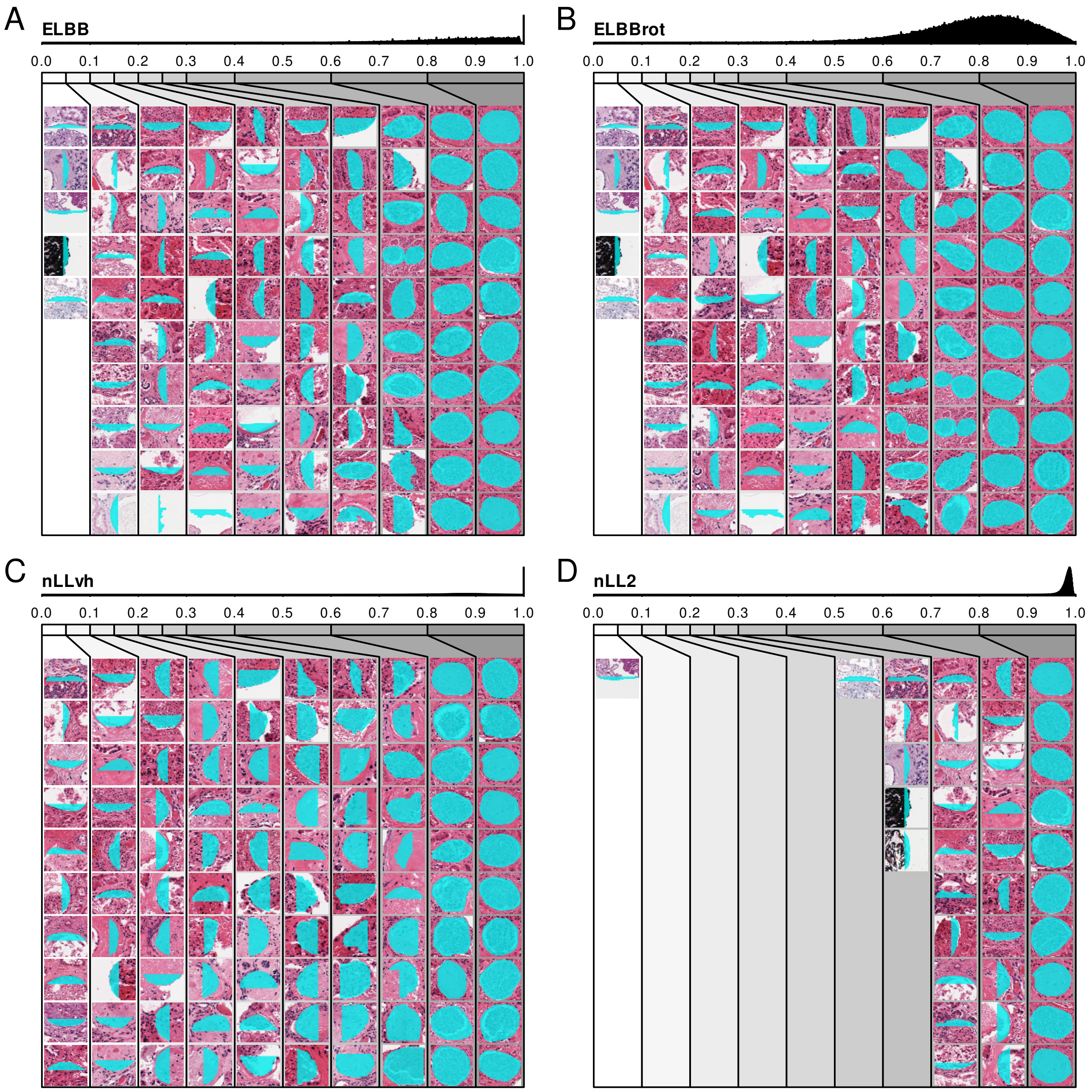

**Supp. Fig. 10.** Visualization of annotations sorted by descriptors based on elongation (A-B) or longest line segments (C-D). The distribution of descriptor values is divided into 10 bins, and up to 10 random annotations are shown from each bin.

#### Supplementary figure 11

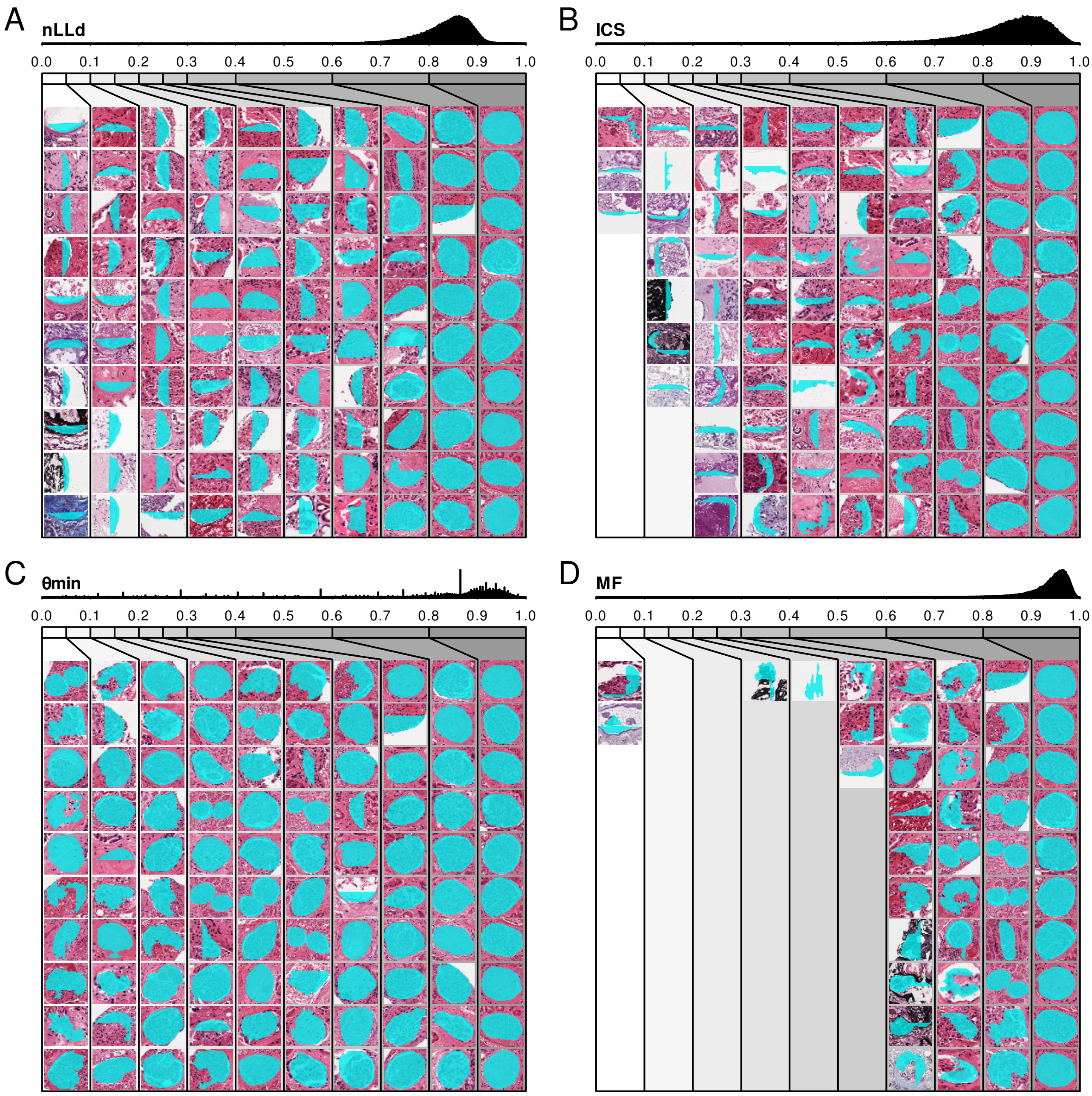

**Supp. Fig. 11.** Visualization of annotations sorted by the descriptors based on longest diagonal line segments (A), inscribed circle sphericity (B), the smallest boundary angle (C), or the Moments-based feature (D). The distribution of descriptor values is divided into 10 bins, and up to 10 random annotations are shown from each bin.

#### Supplementary figure 12

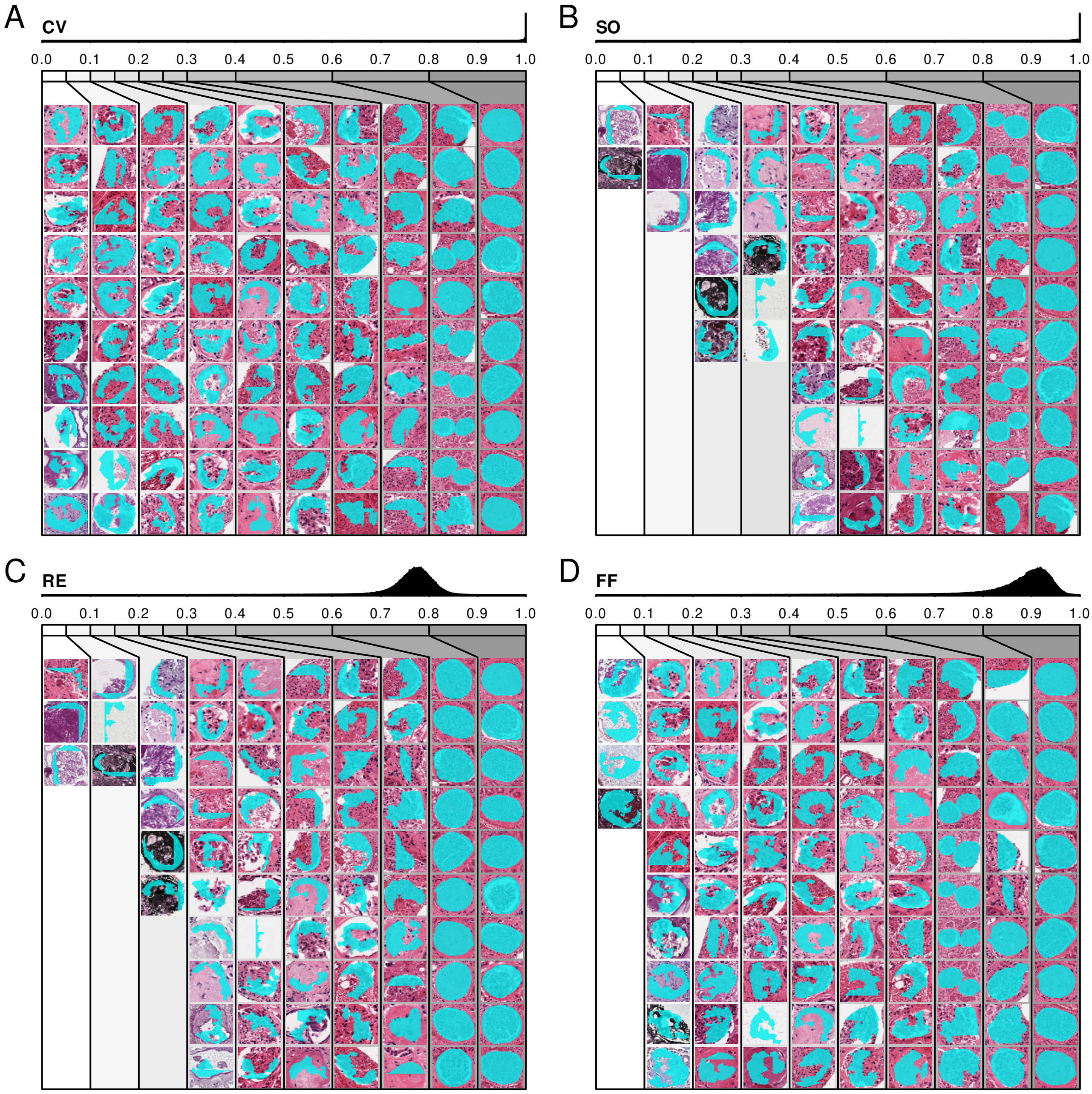

**Supp. Fig. 12.** Visualization of annotations sorted by the descriptors based on convexity (A), solidity (B), rectangularity (C), or the Fourier descriptor (D). The distribution of descriptor values is divided into 10 bins, and up to 10 random annotations are shown from each bin.

#### Supplementary figure 13

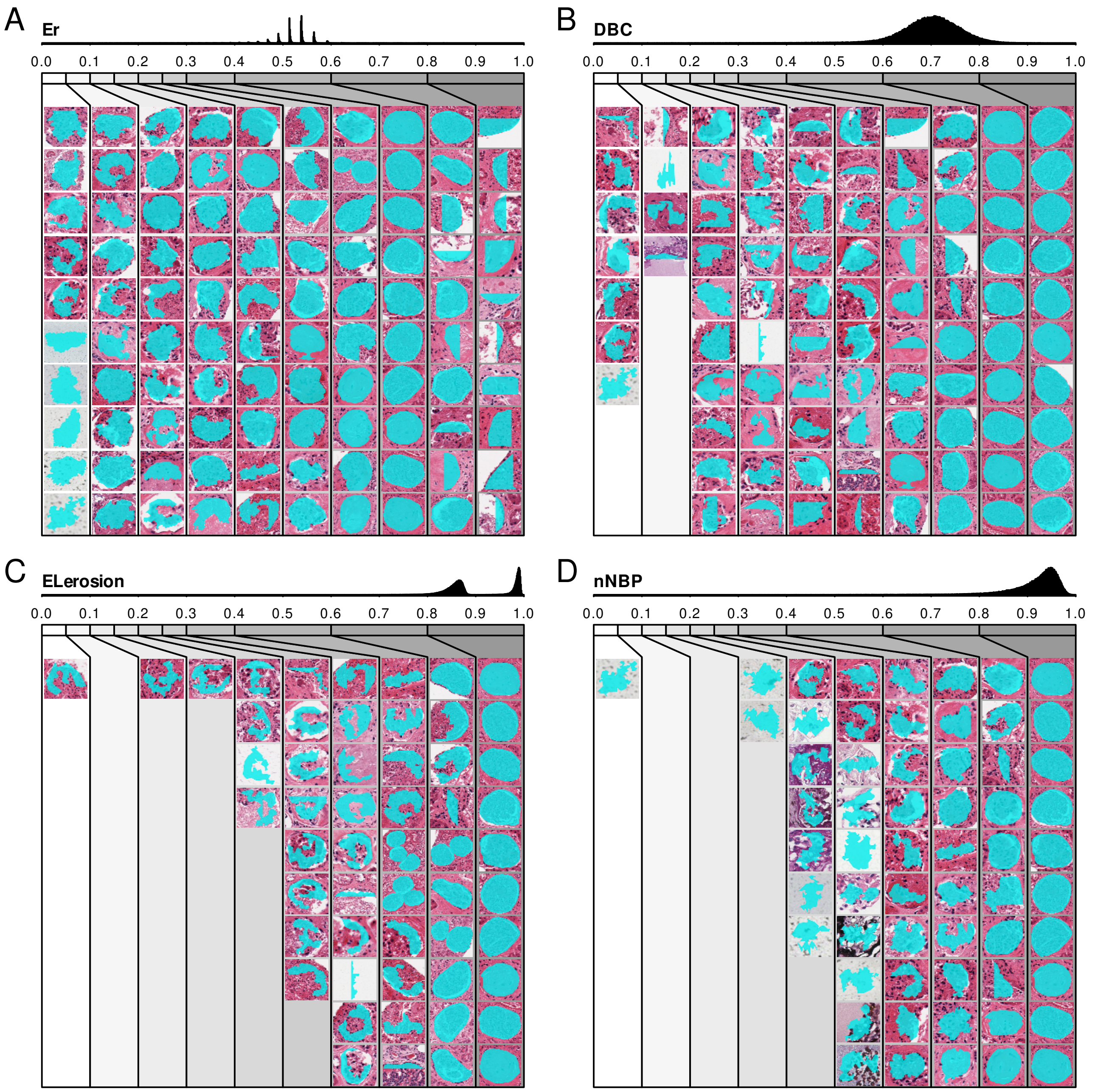

**Supp. Fig. 13.** Visualization of annotations sorted by the descriptors based on the entropy of radial distances (A), the fractal dimension based on box counting (B), the erosion-based elongation (C), or the normalized number of boundary points (D). The distribution of descriptor values is divided into 10 bins, and up to 10 random annotations are shown from each bin.

#### Supplementary figure 14

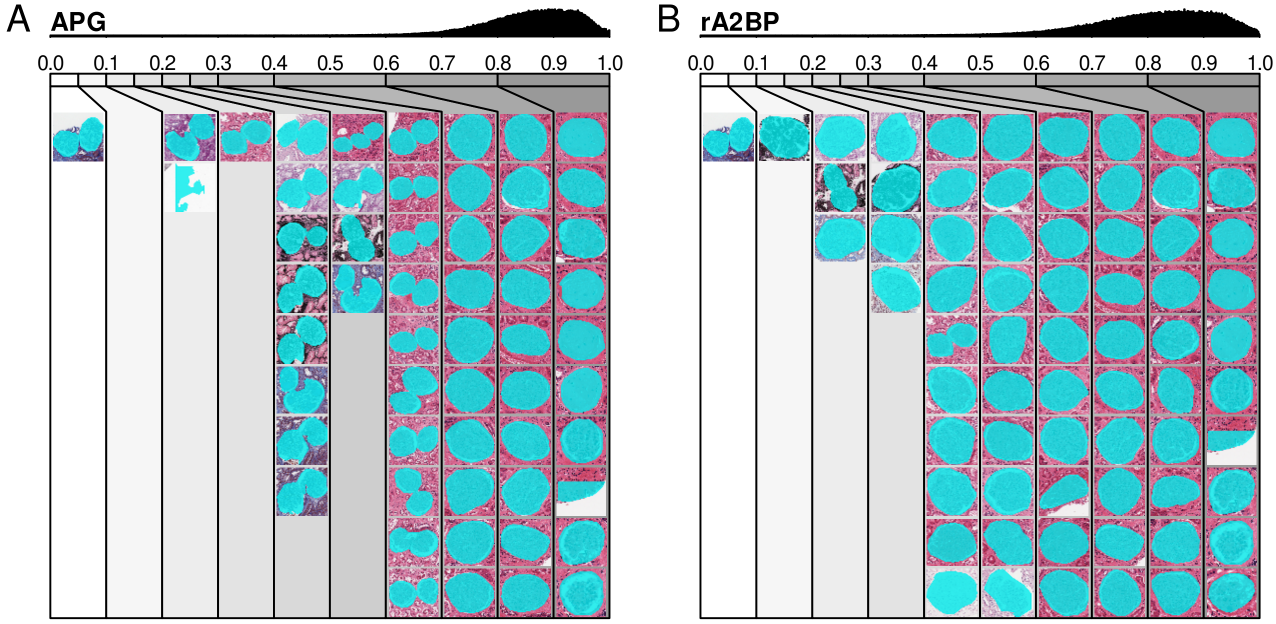

**Supp. Fig. 14.** Visualization of annotations sorted by the descriptors based on polygonal area (A) or the ratio between area and number of boundary points (B). The distribution of descriptor values is divided into 10 bins, and up to 10 random annotations are shown from each bin.

#### Supplementary figure 15

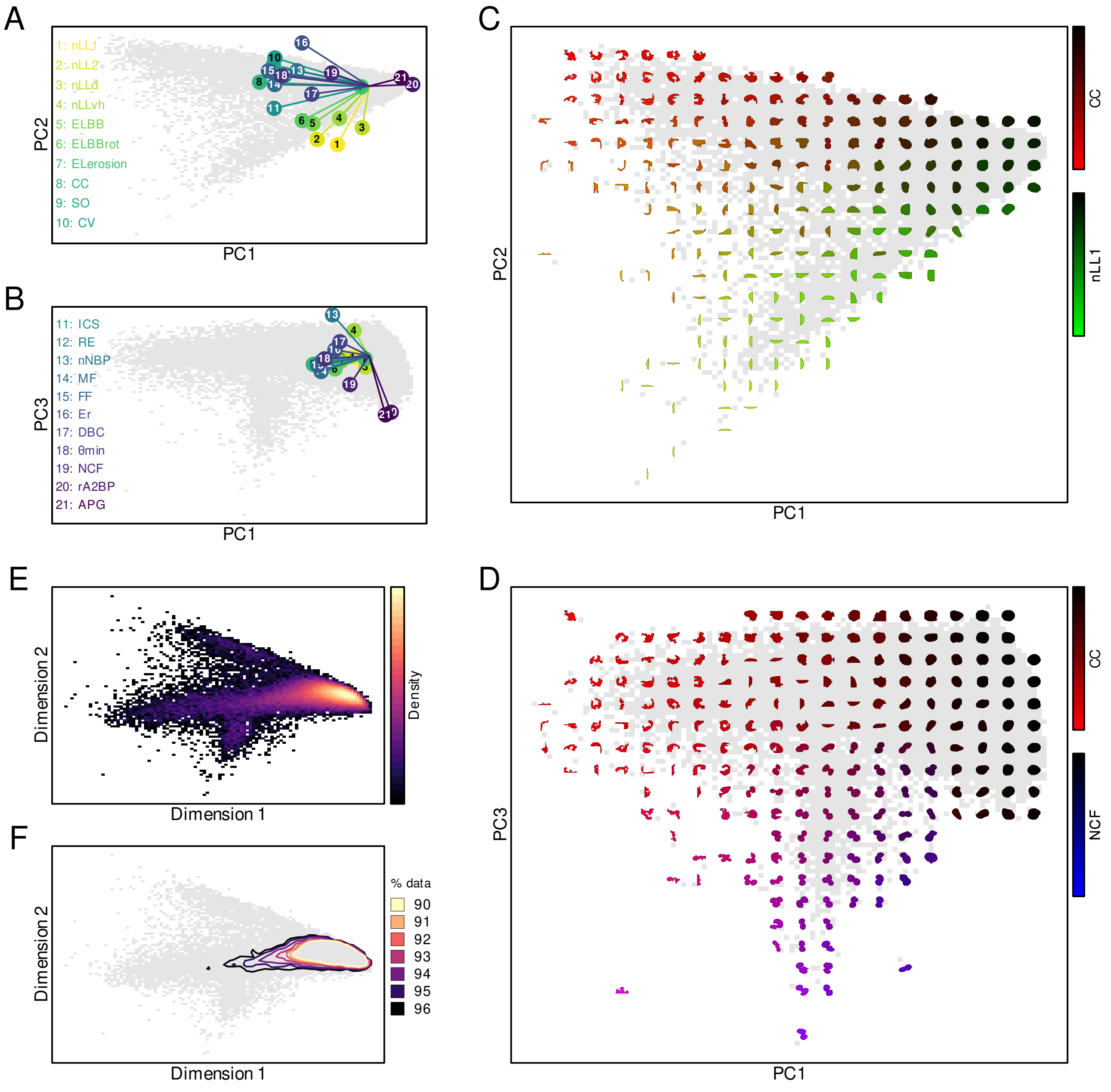

**Supp. Fig. 15.** Analysis of annotations and shape descriptors via PCA and MDS. **A-B)** Biplots showing the loadings of the various descriptors in the two PCA plots comparing the first principal component either to the second (A) or third (B) component. **C-D)** PCA plots comparing the first principal component either to the second (C) or third (D) component, overlaid with example annotations colored based on the values of the *CC* (red, C-D), $nLL_1$ (green, C), and *NCF* (blue, D) descriptors. **E)** Density map of the MDS results showing the distribution of annotations. **F)** Contour plot for the MDS results indicating the regions containing 90%, 91%, 92%, 93%, 94%, 95%, and 96% of the annotations.

#### Supplementary figure 16

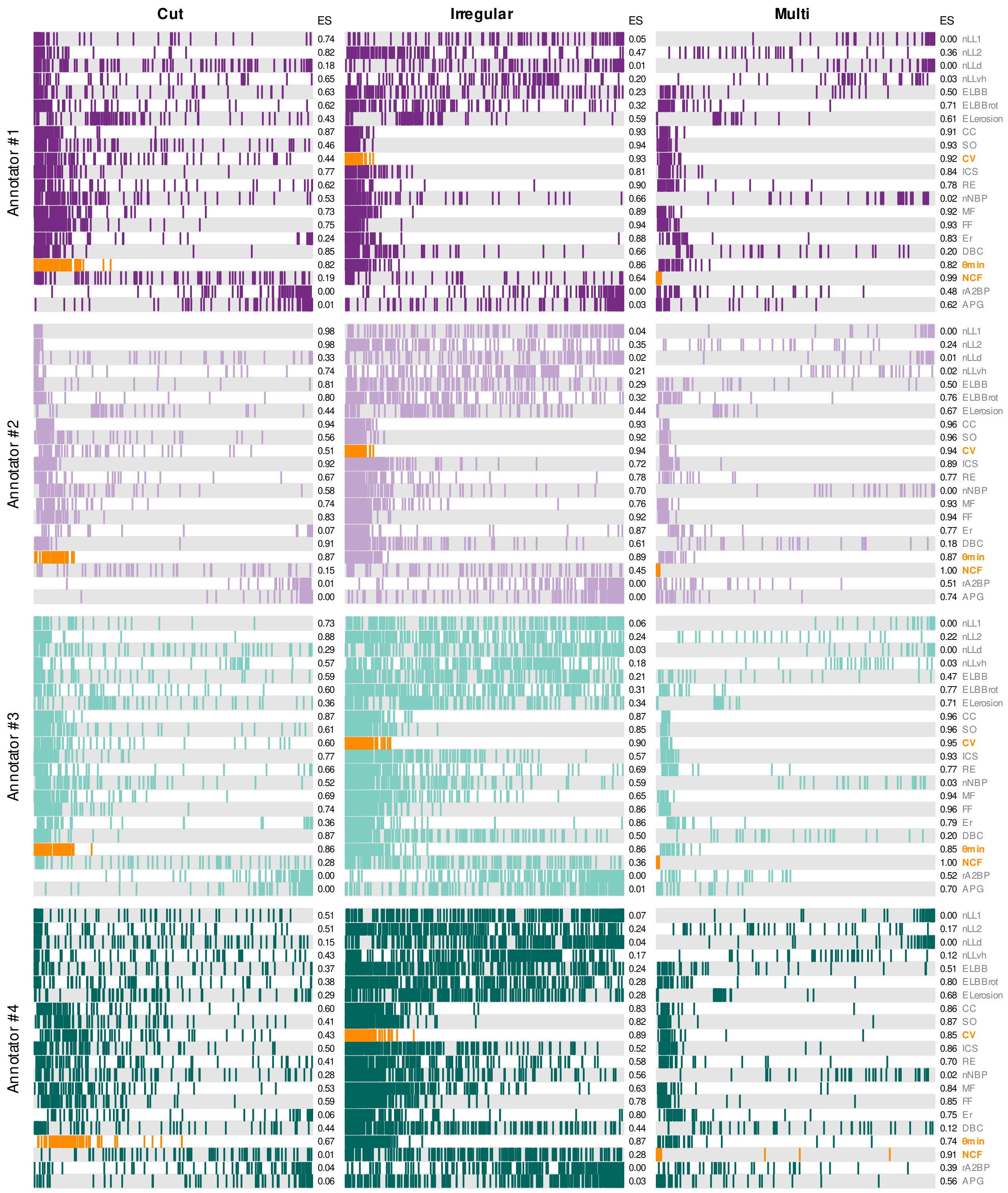

**Supp. Fig. 16.** Enrichment of faulty segmentation annotations by the 21 different shape descriptors. Each column represents one of the three error categories (cut, irregular, multi-object), divided into blocks for each of the four annotators. In each block, a single row visualizes the enrichment of annotations labeled with the respective fault by the respective annotator, after sorting with one of the 21 shape descriptors. Specifically, in a single row, each colored line indicates the position of a fault among the sorted annotations (compare also Supp. fig. 9). Rows highlighted in orange signify the descriptor deemed generally best for enriching the respective error category.

#### Supplementary figure 17

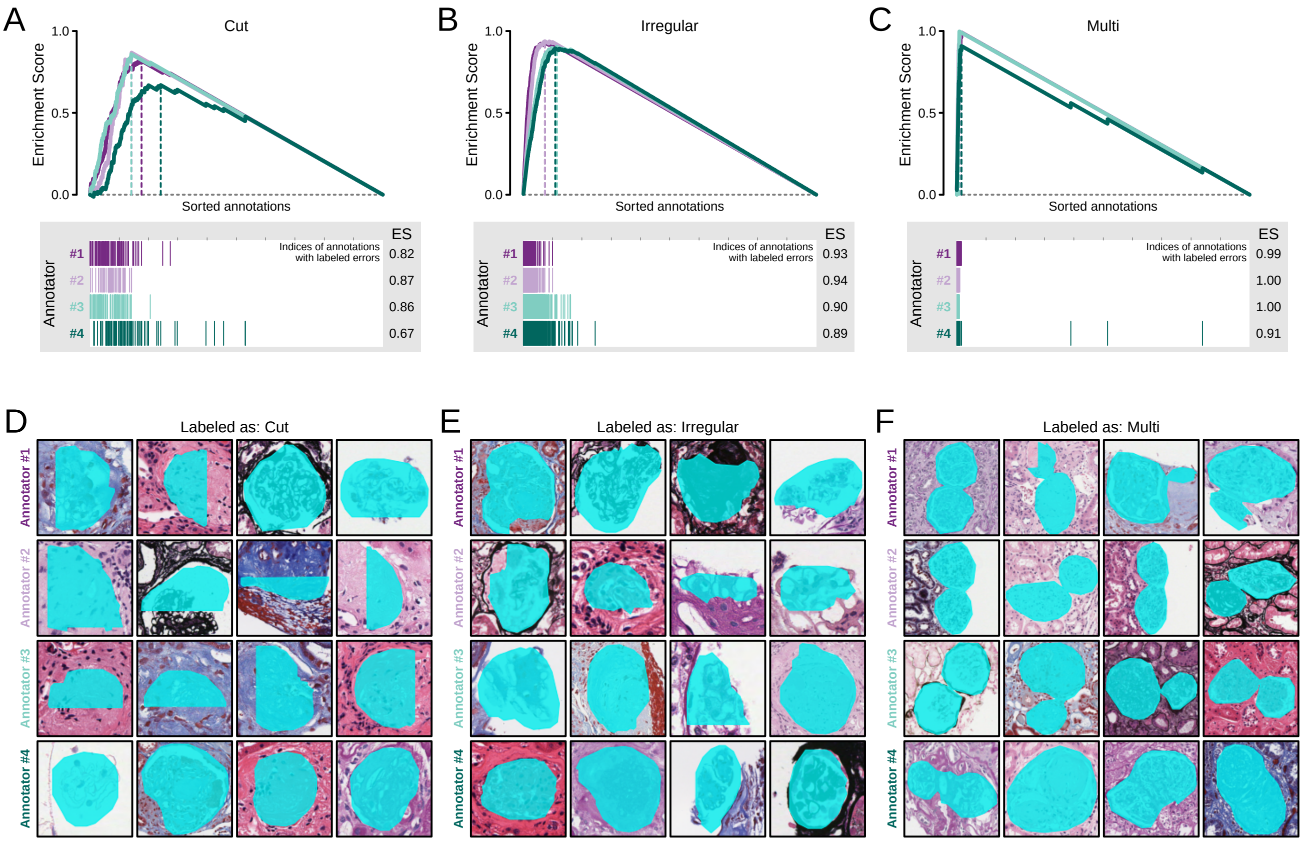

**Supp. Fig. 17.** Quantitative evaluation of shape descriptors. **A-C)** Enrichment analysis of cut (A), irregular (B), and multi-object (C) segmentation faults after sorting annotations using the $\theta_{min}$, $CV$, or $NCF$ descriptor, respectively. The bottom panels display the indices of annotations labeled as faulty (colored lines) among the sorted annotations, and the upper panels display the resulting enrichment curves (See also supp. figure 9). **D-F)** The four highest ranked annotations per category (D: Cut; E: Irregular; F: Multi-object) labeled as faulty by each annotator, respectively.
